# Innovative assembly strategy contributes to the understanding of evolution and conservation genetics of the critically endangered *Solenodon paradoxus* from the island of Hispaniola

**DOI:** 10.1101/164574

**Authors:** Kirill Grigorev, Sergey Kliver, Pavel Dobrynin, Aleksey Komissarov, Walter Wolfsberger, Ksenia Krasheninnikova, Yashira M. Afanador-Hernández, Liz A. Paulino, Rosanna Carreras, Luis E. Rodríguez, Adrell Núñez, Filipe Silva, J. David Hernández-Martich, Audrey J. Majeske, Agostinho Antunes, Alfred L. Roca, Stephen J. O’Brien, Juan Carlos Martinez-Cruzado, Taras K. Oleksyk

## Abstract

Solenodons are insectivores living on the Caribbean islands, with few surviving related taxa. The genus occupies one of the most ancient branches among the placental mammals. The history, unique biology and adaptations of these enigmatic venomous species, can be greatly advanced given the availability of genome data, but the whole genome assembly for solenodons has never been previously performed, partially due to the difficulty in obtaining samples from the field. Island isolation has likely resulted in extreme homozygosity within the Hispaniolan solenodon (*Solenodon paradoxus*), thus we tested the performance of several assembly strategies for performance with genetically impoverished species’ genomes. The string-graph based assembly strategy seems a better choice compared to the conventional de Brujn graph approach, due to the high levels of homozygosity, which is often a hallmark of endemic or endangered species. A consensus reference genome was assembled from sequences of five individuals from the southern subspecies (*S. p. woodi*). In addition, we obtained one additional sequence of the northern subspecies (*S. p. paradoxus*). The resulting genome assemblies were compared to each other, and annotated for genes, with a specific emphasis on the venomous genes, repeats, variable microsatellite loci and other genomic variants. Phylogenetic positioning and selection signatures were inferred based on 4,416 single copy orthologs from 10 other mammals. Patterns of SNP variation allowed us to infer population demography, which indicated a subspecies split within the Hispaniolan solenodon at least 300 Kya.

## Background

The only two surviving species of solenodons found on the two largest Caribbean islands, Hispaniola (*Solenodon paradoxus*) and Cuba (*S. cubanus*) are among the few endemic terrestrial mammals that survived human settlement on these islands. Phenotypically, solenodons resemble shrews (**Figure 1**), but molecular evidence indicates that they are basal to all other eulipotyphlan insectivores, having split from other placental mammals in the Cretaceous Period [1–3]. These enigmatic species have various local names in Cuba and Hispaniola, including *orso* (bear), *hormiguero* (ant-eater), *joron* (ferret), *milqui* (or *almiqui*) and *agouta* [4,5], all pointing to the first impression made on the Spanish colonists by its unusual look. Today, the Hispaniolan solenodon *(Solenodon pardoxus*) is difficult to find in the wild, both because of its nocturnal lifestyle and the low population numbers. Here, we report the assembly and annotation of the nuclear genome sequences and genomic variation of two subspecies of *S. paradoxus*, using analytical strategies that will allow researchers to ask questions and develop tools to assist future studies of evolutionary inference and conservation applications.

**Figure 1.**
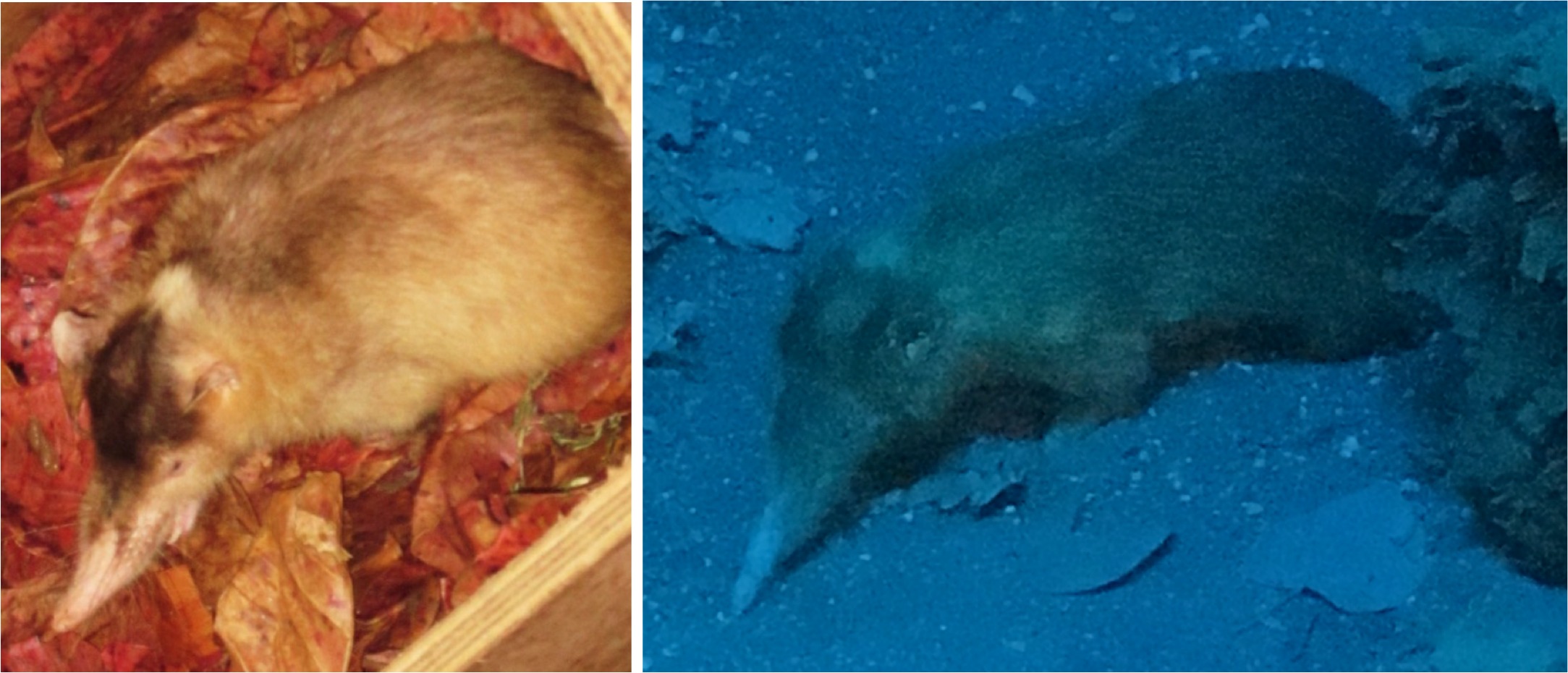
Phenotypic variation. **A)** A captive Hispaniolan solenodon from the northern subspeices (*Solenodon paradoxus paradoxus*) photographed at the Santo Domingo Zoo (photo taken by Juan C. Martinez-Cruzado in 2014). B). A mounted specimen of the southern subspecies (*S. paradoxus woodii*) photographed at the *Museo Nacional de Historia Natural prof. Eugenio de Jesus Marcano* in Santo Domingo, Dominican Republic (photo taken by Taras K. Oleksyk in 2017).

*S. paradoxus* was originally described from a skin and an imperfect skull at the St. Petersburg Academy of Sciences in Russia [6]. It has a large head with a long rostrum with tiny eyes and ears partially hidden by the dusky brown body fur that turns reddish on the sides of the head, throat and upper chest. The tail, legs, snout, and eyelids of the *S. paradoxus* are hairless. The front legs are noticeably more developed, but all four have strong claws, probably useful for digging (**Figure 1**). Adult animals measure 49-72 cm in total length, and weigh almost 1kg [7]. Solenodons are social animals, they spend their days in extensive underground tunnel networks shared with other members of family groups, and come to the surface at night to hunt small vertebrates and large invertebrates [8]. A unique feature is the *os proboscidis*, a bone extending forward from the nasal opening to support the snout cartilage [9]. Solenodons are venomous mammals, that display a fascinating strategy for venom delivery. The second lower incisor of solenodons has a narrow, almost fully enfolded tubular channel, through which saliva secreted by the submaxillary gland flows into the victim (Folinsbee et al. 2007). (The genus name, “solenodon,” means “grooved tooth” in Greek and refers to the shape of this incisor). Although solenodons rarely bite humans, the bites can be very painful (Nicolás Corona, personal communication), and even a small injection of venom has been shown to be fatal to mice in minutes [7]. The chemical composition of the solenodon venom has not yet been resolved [10].

Morphometric studies suggest that southern and northern solenodons may be distinctive enough to be considered separate subspecies [2,11,12], a notion supported by recent mitochondrial DNA studies [13,14]. Roca et al. 2004 sequenced relatively short mitochondrial fragments spanning 2.5-kilobase portions of both mitochondrial ribosomal RNA genes of *S. paradoxus* and the Cuban solenodon (*S. cubanus*), implying that solenodon divergence from other eulipotyphlan mammals such as shrews and moles date back to the Cretaceous era, ~76 million years ago (Mya), before the mass extinction of the dinosaurs ~ 65 Mya. Brandt et al. 2016 sequenced complete mitogenome sequences of six Hispaniolan solenodon specimens, corroborating this conclusion, and estimated that *S. paradoxus* diverged from all other mammals approximately 76 Mya. An analysis of five nuclear genes gave a much later estimate (<60 Mya) for the solenodon divergence [15], and disagreed on the date and the mode of speciation of the two extant species. Specifically, this study suggested a much more recent date for speciation following a Cenozoic over-water dispersal 3.7–4.8 Mya [15], rather than vicariance following land separation between Eastern Cuba and Western Hispaniola 25 Mya [3]. Current molecular data allows the whole genome analysis of *S. paradoxus* that can provide support and validation to the earlier evolutionary studies.

It may now be imperative to study conservation genomics of solenodons, whose extinction would extirpate an entire evolutionary lineage whose antiquity goes back to the age of dinosaurs. *S. paradoxus* survived in spectacular island isolation despite the devastating human impact to biodiversity in recent centuries [3,13]. Nevertheless, survival of this species is now threatened by deforestation, increasing human activity, and predation by introduced dogs, cats and mongooses, and it is listed as endangered (B2ab, accessed in 2008), with its habitat severely fragmented and declining in population by the IUCN Red List of Threatened Species (http://www.iucnredlist.org/details/20321/0).

In this study, we assembled the genome of *S. paradoxus* using low coverage genome data (~5x each) from five *S. paradoxus woodi* individuals. We take advantage of the low individual and population genetic diversity to pool individual data, and apply a string graph assembly approach resulting in a working genome assembly of the *S. paradoxus* genome from the combined paired-end dataset (approximately 26x). Our methodology introduces a useful pipeline for genome assembly to compensate for the limited amount of sequencing, which, in this instance performs better than the assembly by a traditional de Bruijn algorithm (SOAPdenovo2) [16]. We employed the string-graph assembler Fermi [17] as a principal tool for contig assembly in conjunction with SSPACE [18] and GapCloser [16] for scaffolding. The resulting genome sequence data was sufficient for high-quality annotation of genes and functional elements, as well as for comparative genomics and population genetic analyses. Prior to this study, the string-graph assembler Fermi [17] has been used only in studies for annotation, or as a complementary tool for *de novo* assemblies made with de Bruijn algorithms [19]. We present and compare genome assemblies for the southern subspecies (*S. p. woodi*) based on several combinations of assembly tools, provide a high-quality annotation of genome features and describe genetic variation in two subspecies (*S. p. woodi and S. p. paradoxus*), make inferences about recent evolution and selection signatures in genes, trace demographic histories, and develop molecular tools for future conservation studies.

## Data description

### Sample collection and sequencing

Five *S. paradoxus woodi* adult individuals from southern Dominican Republic were collected in the wild following the general field protocol including two specimens caught from La Cañada del Verraco, and three from the El Manguito location in the Pedernales Province. In addition, one *S. p. paradoxus* (Spa-1) sample was acquired through the collaboration with ZooDom at Santo Domingo, but was originally brought there from Cordillera Septentrional in the northern part of the island. The captured individuals were visually assessed for obvious signs of disease, weighed, measured, sexed, and released at the capture site, all within 10 minutes of capture. Geographic coordinates were recorded for every location. **Figure 2** highlights geographical locations of sample collection points for the samples used in this study.

**Figure 2.**
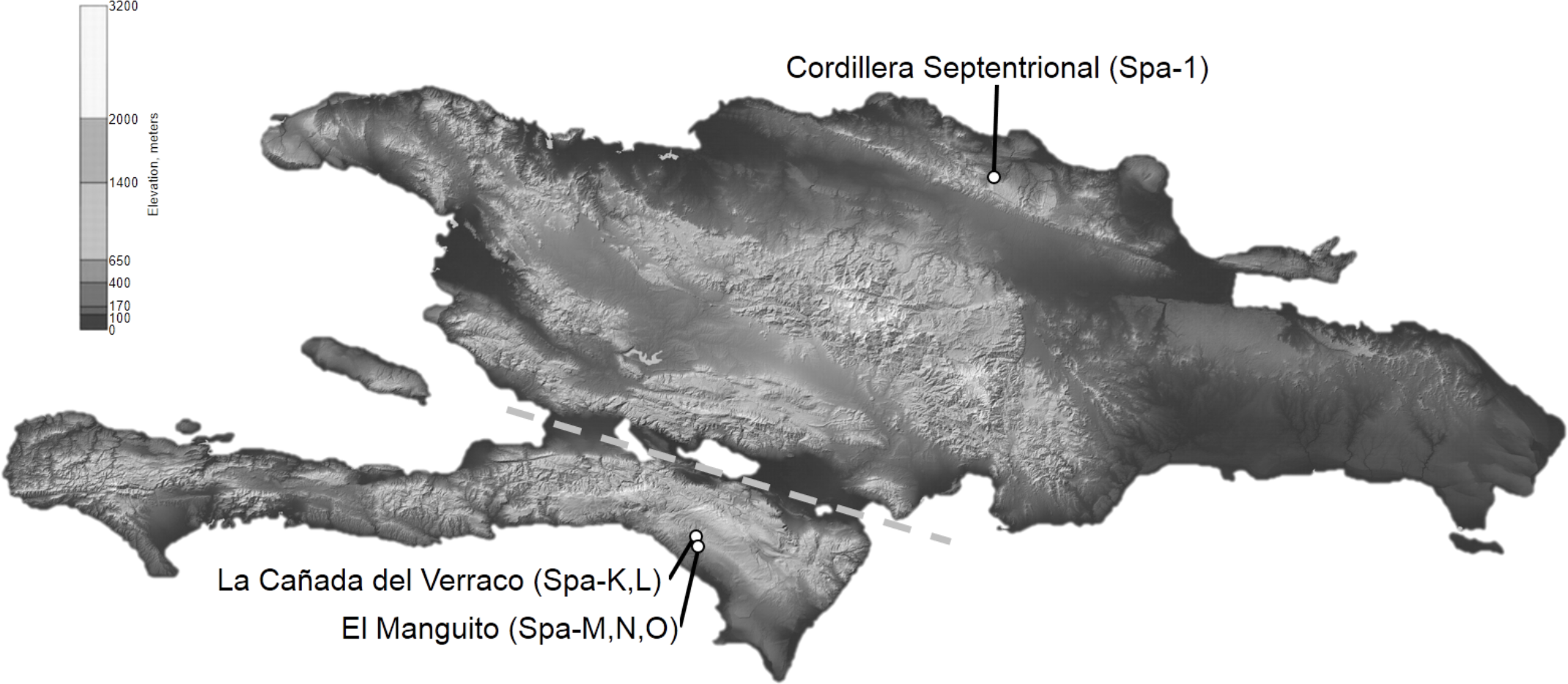
Origins of the genomic DNA samples of *Solenodon paradoxus* from the island of Hispaniola. Approximate locations of capture for five wild individuals of *S. p. woodi*: Spa-K and Spa-L from La Cañada del Verraco, as well as Spa-M, Spa-N, and Spa-O from the El Manguito location in the Pedernales Province in the southwest corner of the Dominican Republic bordering Haiti. In addition, one *S. p. paradoxus* sample (Spa-1) from Cordillera Septentrional in the northern part of the island. Exact coordinates of each sample location are listed in Brand et al. 2016. The dashed line indicates the position of the Cul de Sac Plain and Neiba Valley; this region was periodically inundated by a marine canal that separated Hispaniola into north and south paleoislands during the Pliocene and Pleistocene (Ottenwalder 2001). The original map is in the public domain (courtesy of NASA), and is modified from Brandt et al. 2016.

The five *S. p. woodi* samples were sequenced using Hiseq2000 technology (Illumina Inc.), resulting in an average of 151,783,327 paired-end reads, or 15.33Gb of sequence data, per individual. In addition, DNA extracted from the northern solenodon (*S. p. paradoxus*) Spa-1 produced a total of 52,358,830 paired-end reads, equating to approximately 13.09Gb of sequence data. Only the samples of *S. paradoxus woodi* were used for assembly since the northern subspecies (*S. paradoxus paradoxus*) did not have sufficient coverage for the *de novo* assembly.

Further details about sample collection, DNA extraction, library construction and sequencing can be found in the Methods section. The whole genome shotgun data from this project has been deposited at DDBJ/ENA/GenBank under the accession NKTL00000000. The version described in this paper is version NKTL01000000. The genome data has also been deposited into NCBI under BioProject PRJNA368679, and to GigaDB (Grigorev et al. 2017).

### Read correction

After the reduction of adapter contamination with Cookiecutter [20], the k-mer distribution in the reads for the five individuals of *S. paradoxus woodi* was assessed with Jellyfish [21]. The predicted mean genome coverage was approximately 5x for each sample (**Figure 3**). Given the hypothesized low levels of genetic diversity, and in order to increase the average depth of coverage, the reads from the five samples were combined into a single data set. As a result, the projected mean genome coverage for the combined genome assembly was 26x. Error correction was applied with QuorUM [22] using the value k = 31. The k-mer distribution analysis by Jellyfish in the combined and error-corrected data set indicated very low levels of heterozygosity in accordance with the hypothesis (see **Figure 3** legend), allowing use of the combined dataset for the further genome assembly. The genome size has been estimated using KmerGenie [23] to be 2.06Gbp.

**Figure 3.**
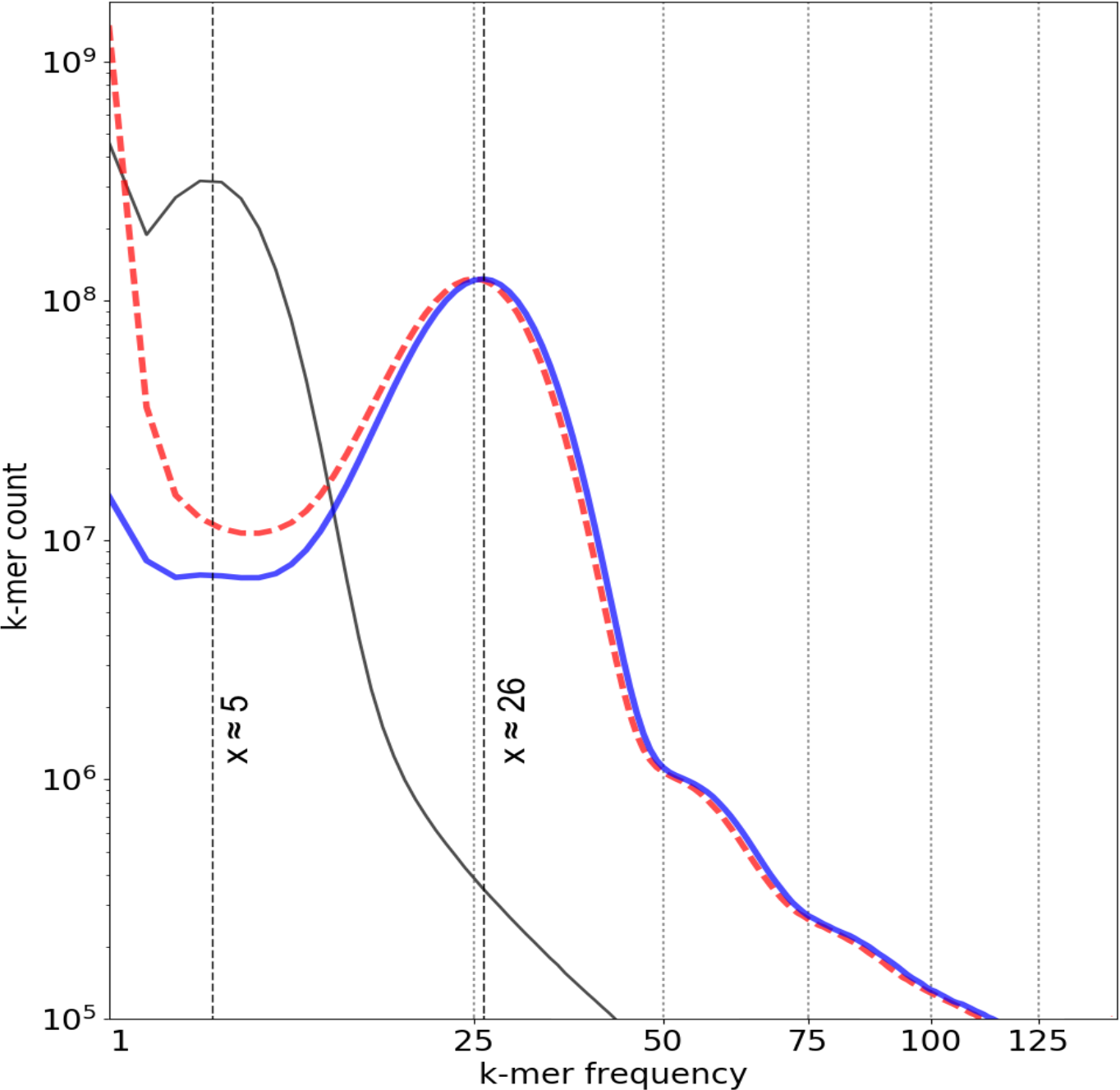
Heterozygosity and k-mer distribution. k-mer distributions for the *S. p. woodi* reads. Only one original sample (SPA-K) distribution is shown with a solid black line as they are identical for the original samples. The predicted mean genome coverage was approximately 5x for each sample (x=5). The combined uncorrected dataset is plotted in a dashed red line. The combined dataset corrected with QuorUM (Marçais et al. 2015) is plotted in a solid blue line. Local maximum on the left-hand side for each distribution (representing k-mers found once or very few times) indicates contribution of sequencing errors. The largest local maxima(to the right) are interpreted as projected coverage. For the combined sample this value is x=26. Smaller local maxima are interpreted as heterozygous contribution; it proves insignificant in the combined sample even after read correction.

## Analyses

### Assembly tool combinations

We used several alternative combinations of tools to determine the best approach to an assembly of the combined genome data, outlined in **Table 1**. First, the combined libraries of paired end reads were assembled into contigs with Fermi, a string graph based tool [17]. Second, the same libraries were also assembled with SOAPdenovo2, a de Bruijn graph based tool [16]. The optimal kmer length parameter for SOAPdenovo2 was determined to be k = 35 with the use of KmerGenie [23]. For the scaffolding step we used either SSPACE [18] or the scaffolding module of SOAPdenovo2 [16]. Finally, for all instances, the GapCloser module of SOAPdenovo2 was used to fill in gaps in the scaffolds [16]. After assembly, datasets were trimmed: scaffolds shorter than 1Kbp were removed from the output. In **Table 1**, the four possible combinations of tools used for the assembly are referred to with capital letters A, B, C, and D for brevity. However, SOAPdenovo2 introduces artifacts at the contig construction stage, which it is specifically designed to mitigate at later stages, and SSPACE is not aware of such artifacts [24]. For this reason, the assembly produced by combination D (contig assembly with SOAPdenovo2 and scaffolding with SSPACE) was not reported.

**Table 1.** Description of the assembly strategies and comparison of metrics for the resulting assemblies

| Assembly Names | A | B | C | D |
| --- | --- | --- | --- | --- |
| **Assembly Tools** |  |  |  |  |
| Contig assembly tool | *Fermi* | *Fermi* | *SOAPdenovo2* | *SOAPdenovo2* |
| Scaffolding tool | *SOAPdenovo2* | *SSPACE* | *SOAPdenovo2* | *SSPACE* |
| Gap closing tool | *GapCloser* | *GapCloser* | *GapCloser* | *GapCloser* |
| **Assembly Metrics** |  |  |  |  |
| Total contigs (>1,000 bp) | 71,429 | 71,429 | 189,566 | 189,566 |
| Contig N50 | 54,944 | 54,944 | 4,048 | 4,048 |
| Contig CEGMA (%) * | 96.37(77.42) | 96.37(77.42) | 68.15(33.06) | 68.15(33.06) |
| Contig BUSCO (%) | 86(65) | 86(65) | 42(21) | 42(21) |
| Total scaffolds (>1,000 bp) | 14,417 | 40,372 | 20,466 | - |
| Final N50 | 555,585 | 110,915 | 331,639 | - |
| Final CEGMA (%) | 95.56(81.85) | 95.97(88.71) | 95.97(90.73) | - |
| Final BUSCO (%) | 91(74) | 86(64) | 94(80) | - |
| Percentage of Ns (%) | 0.06322 | 0.0135 | 0.02622 | - |
| REAPR error-free bases (%) | 96.46 | 95.35 | 94.98 | - |
| REAPR low-scoring regions | 18 | 16 | 71 | - |
| REAPR incorrectly oriented reads | 11,543 | 5,329 | 28,964 | - |

\* BUSCO [26] and CEGMA [27] percentages are reported for all genes (complete and partial), while the percentage of complete genes are shown in parentheses.

### QC and structural comparisons between the assemblies

We used QUAST [25] to estimate the common metrics of assembly quality for all combinations of assembly tools: N50 and gappedness (the percentage of *N*s (**Table 1)**). Fermi assembled contigs (A and B) were overall longer and fewer in number than the SOAPdenovo2 (C and D). The assembly completeness was also evaluated with both BUSCO [26] and CEGMA [27] for completeness of conservative genes. Fermi assemblies (A and B) showed high levels of completeness compared to SOAPdenovo2 (86% vs 42%) at the contig level. However, this difference is partially mitigated at the scaffolding step where SOAPdenovo2 increases completeness for Fermi assembly (A), and more than doubles it for the SOAPdenovo2 assembly (C). To directly evaluate the quality of all the assemblies we applied REAPR [28]. From the REAPR metrics presented at the bottom part of Table 1, it appears that, even though the scaffolding step has increased the final N50 for the C assembly, it contains significantly more regions with high probability of misassemblies (low-scoring regions), less error-free bases, and 3 to 6 times higher number of incorrectly oriented reads compared to the Fermi based assemblies (A and B) (**Table 1**).

We hypothesized that aligning the three genome assemblies to each other will allow us to detect some of these misassembles. A comparison to the best, most closely related genome assembly (e.g. *Sorex araneus*) will reveal several rearrangements that in many cases reflect real evolutionary events. It is reasonable to assume that, if all the rearrangements that are detected are real, and not due to the assembly artifacts, the number of detected rearrangements vs *Sorex* assembly should be the same for all three *Solenodon* assemblies (A, B and C). Following the parsimony principle, an assembly showing rearrangements is also likely to be containing the most assembly artifacts. Conversely, we expected that the best of the three assemblies of the *Solenodon* genome should contain the least number of reversals and transpositions when compared to the best available closely related genome (*Sorex araneus*).

To test this hypothesis, the three completed assemblies of *Solenodon* (A, B and C) were aligned to each other, and to the outgroup, which was the *Sorex* genome (SorAra 2.0, NCBI accession number GCA_000181275.2), using Progressive Cactus [29]. Custom scripts were employed to interpret binary output of the pairwise genome by genome comparisons, and the resulting coverage metrics are presented in **Table 2**. In this comparison, all three *Solenodon* genome assemblies had a significant overlap, and resulted in similar levels of synteny when compared against the *Sorex* reference assembly, but assemblies A and B were the most closely related, while assembly C was slightly more different from each of them. Next, syntenic blocks between each of the three *Solenodon* assemblies (A, B and C) were compared to the *Sorex* assembly, and 50Kbp syntenic blocks were identified using the ragout-maf2 synteny module of the software package *Ragout* [30], and the numbers of scaffolds that contained syntenic block rearrangements were determined. As a result, assembly B had the lowest number of reversals and transpositions when compared to the *S. araneus* reference genome (**Table 2**). Based on the combined results of the evaluations by REAPR [28], Progressive Cactus [29] and Ragout [30], assembly C (generated by the complete SOAPdenovo2 run) was not included in the further analysis.

**Table 2.**
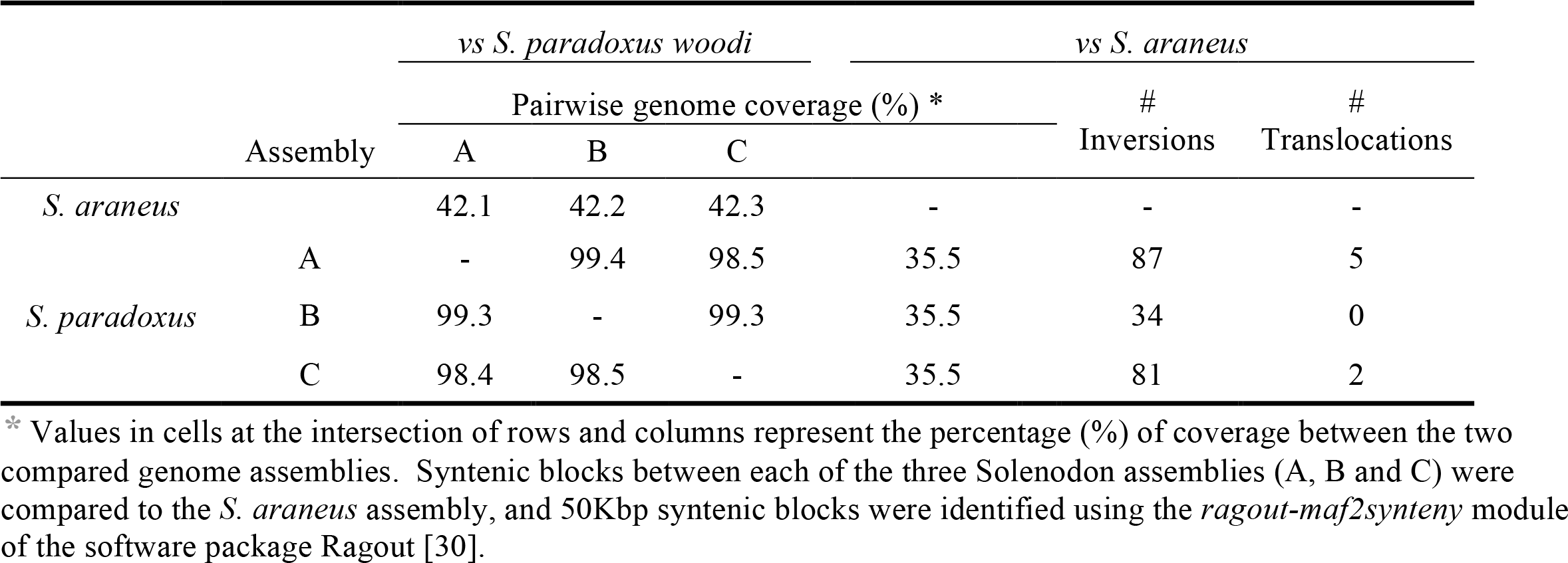
Pairwise genomic coverage for the three assemblies and the *Sorex araneus* genome (SorAra 2.0, NCBI accession number GCA_000181275.2) obtained from the *Progressive Cactus* [29] alignments. While all three assemblies have similar amounts of syntenic coverage to the *Sorex* genome, assembly B contains the least numbers of structural rearrangements (inversional and translocations) compared to the other two assemblies (A and C).

### Genome annotation and evaluation of assembly completeness

Repeats in assemblies A and B were identified and soft masked using RepeatMasker [31] with the RepBase library [32]. The total percentage of all interspersed repeats masked in the genome was lower than in *S. araneus* (22.53% vs 30.48%). This occurred maybe because a low coverage assembly was likely to perform better in non-repetitive regions. Alternatively, if the repeat content in *S. paradoxus* is indeed lower, it will have to be evaluated using a higher quality assembly with the use of long read data. The total masked repeat content of the *S. paradoxus* genome including simple/tandem repeats, satellite DNA, and low complexity regions, etc. is presented in **Table 3**. The repeat content can be retrieved from **Database S1**.

**Table 3.**
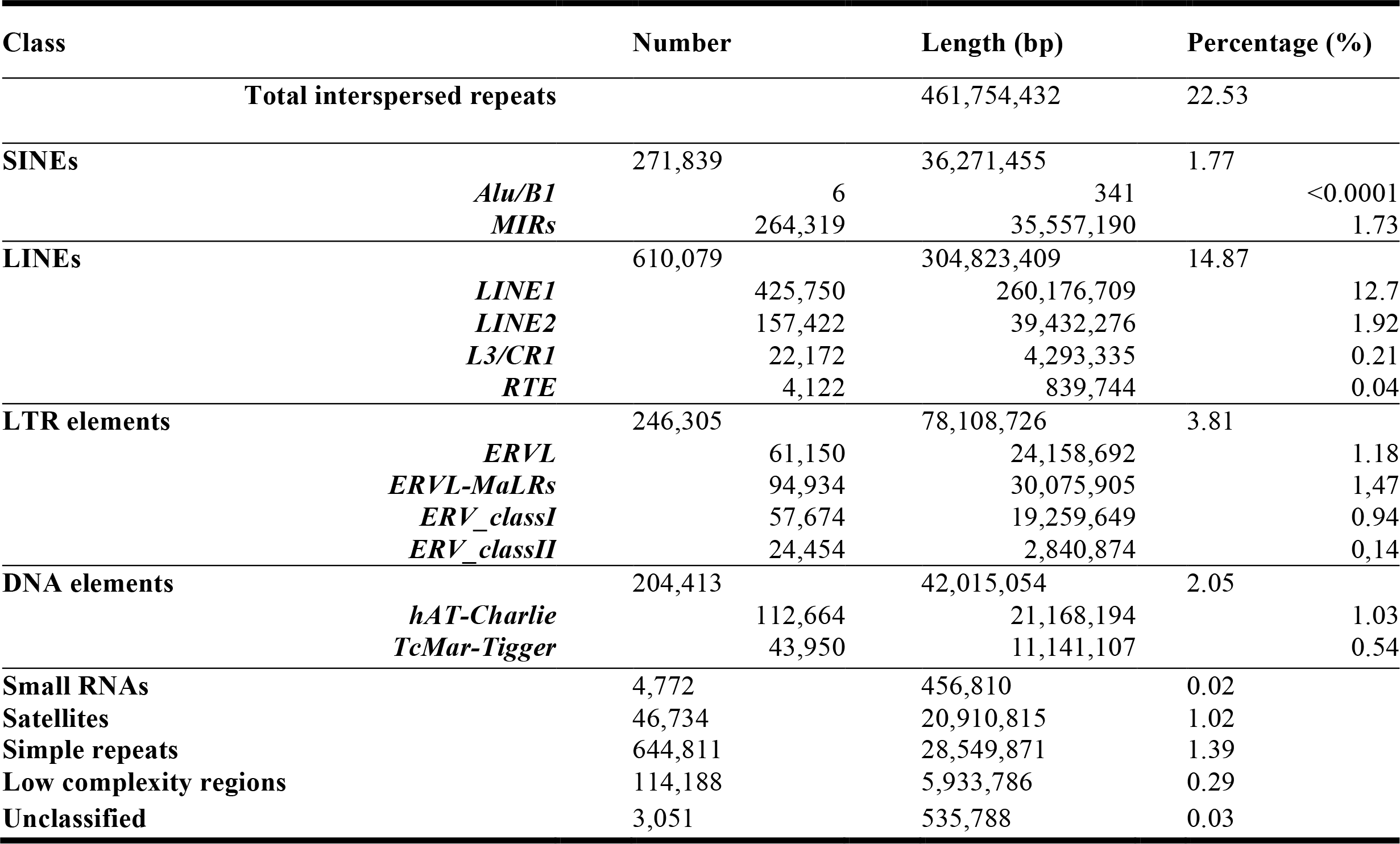
Repeat content of the *Solenodon paradoxus* genome (Assembly B), annotated by RepeatMasker [31] with the RepBase library [32].

The annotation of protein-coding genes was performed using a combined approach that synthesized both homology-based and *de novo* predictions, where *de novo* predictions were used to fill gaps and extend homology-based predictions. Gene annotation was performed for both assemblies (A and B) independently. Proteins of four reference species *S. araneus* (SorAra 2.0, GCA_000181275.2), *Erinaceus europaeus* (EriEur2.0, GCA_000296755.1), *Homo sapiens* (GRCh38.p7) and *Mus musculus* (GRCm38.p4) were aligned to a *S. paradoxus* assembly with Exonerate [33] with a maximum of three hits per protein. The obtained alignments were classified into top (primary) and secondary; the CDS fragments were cut from each side by 3bp for the top hits and by 9bp for secondary hits. These truncated fragments were clustered and supplied as *hints* (local pieces of information about the gene in the input sequence, such as a likely stretch of coding sequence) of the potentially protein-coding regions to the AUGUSTUS software package [34], which predicted genes in the soft-masked *Solenodon* assembly. Proteins were extracted from the predicted genes and aligned by HMMER [35] and BLAST [36] to Pfam [37] and Swiss-Prot (UniProt Consortium & others, 2014) databases, respectively. Genes supported by hits to protein databases and hints were retained; the unsupported sequences were discarded. The annotated genes can be retrieved from **Database S2**.

Assembly B showed a higher support compared to assembly A (91.7% vs 79.2%) for the protein coding gene predictions by extrinsic evidence, even though assembly A had a larger N50 value (**Table 1**). These values were calculated as a median fraction of exons supported by alignments of proteins from reference species to genome (**Figure 4**). In other words, assembly B is more useful for gene predictions, and is likely to contain better gene models that can be used in the downstream analysis. Therefore, based on two lines of evidence: low rearrangement counts (**Table 2**), and high support to gene prediction for the assembly B, it was chosen for the subsequent analyses as the most useful current representation of the *Solenodon* genome.

**Figure 4.**
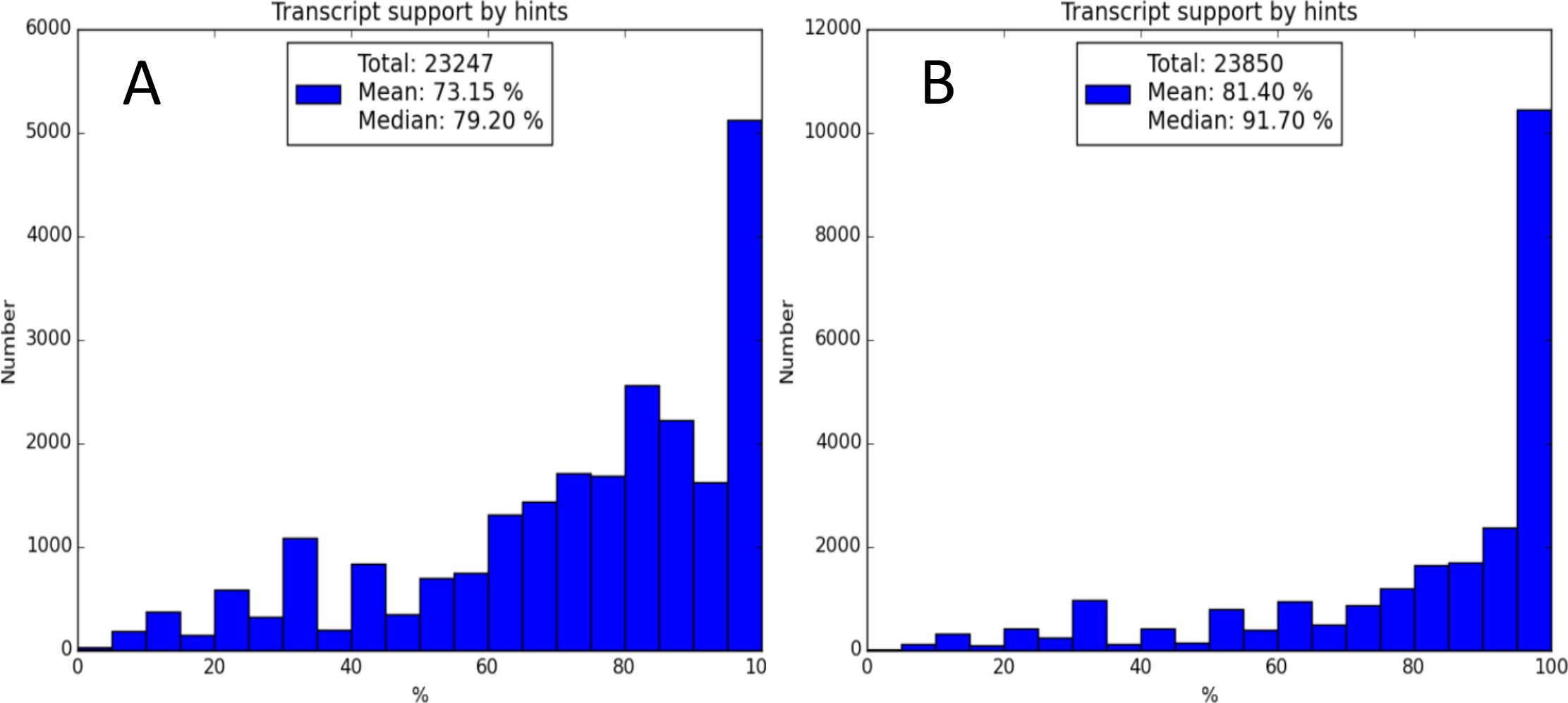
Distribution of the gene prediction support by extrinsic evidence for *Solenodon* assemblies A (on the left) and B (on the right). Proteins of four reference species *S. araneus* (SorAra 2.0, GCA_000181275.2), *Erinaceus europaeus* (EriEur2.0, GCA_000296755.1), *Homo sapiens* (GRCh38.p7) and *Mus musculus* (GRCm38.p4) were aligned to a *S. paradoxus* assembly with Exonerate [33] with a maximum of three hits per protein. Coding sequences (CDS) were cut from each, clustered and uploaded into the AUGUSTUS software package [34] to predict genes in the soft-masked *Solenodon* assembly. Proteins from the predicted genes were aligned by HMMER [35] and BLAST [36] to Pfam [37] and Swiss-Prot [38] databases. Genes supported by hits to protein databases and hints were retained; the rest were discarded. Significantly more transcripts have higher hint support in assembly B. The annotated genes can be retrieved from **Database S2**. Assembly C has not been evaluated.

### Non-coding RNA genes

For all non-coding RNA genes except for tRNA and rRNA genes, the search was performed with INFERNAL (Nawrocki and Eddy 2013) using the Rfam [39] BLASTN hits as seeds. The tRNA genes were predicted using tRNAScan-SE [40], and rRNA genes were predicted with Barrnap ((*BAsic Rapid Ribosomal RNA Predictor*) version 0.6 [41]). Additionally, RNA genes discovered by RepeatMasker at the earlier stages of the analysis were used to cross-reference the findings of rRNA and tRNA-finding software. The list of the non-coding RNA genes can be accessed in **Database S3**.

### Multiple genome alignment, synteny and duplication structure

To compare the duplication structure of the *Solenodon* genome assembly with other mammalian genomes, a multiple alignment with genomes of related species was performed using Progressive Cactus [29]. Currently available genomic assemblies of cow (*Bos taurus*, BosTau 3.1.1, NCBI accession number DAAA00000000.2), dog (*Canis familiaris*, CanFam 3.1, GCA_000002285.2), star nosed mole (*Condylura cristata*, ConCri 1.0, GCF_000260355.1), common shrew (*S. araneus*, SorAra 2.0, GCA_000181275.2) and *S. paradoxus woodi* (assembly B from this study) were aligned together, guided by a cladogram representing branching order in a subset of a larger phylogeny (**Figure S1**). We evaluated the *S. paradoxus* coverage by comparing it to the weighted coverages of other genomes in the alignment to the *C. familiaris* genome (**Table 4**). Custom scripts were employed to interpret the binary output of Progressive Cactus [29]. Cactus genome alignments were used to build a “sparse map’” of the homologies between a set of input sequences. Once this sparse map is constructed, in the form of a Cactus graph, the sequences that were initially unaligned in the sparse map are also aligned [29]. Weighted coverage of a genome by a genome was calculated by binning an alignment into regions of different coverage and averaging these coverages, with lengths of bins as weights. The weighted coverage of *S. paradoxus* to *C. familiaris* was 1.05, which indicated that the present genome assembly is comparable in quality and duplication structure to other available mammalian assemblies, which are close to each other and are close to 1.0 (**Table 4**).

**Table 4.** The weighted coverages of the genomes in the Progressive Cactus alignment [29], as calculated against the *C. familiaris* genome. The weighted coverage of the *S. paradoxus* genome assembly from our study is comparable to other high coverage mammalian genome assemblies. The cladogram used for multiple genome alignment with Progressive Cactus is shown in **Figure S1**.

| Query genome | Weighted<br>coverage |
| --- | --- |
| Dog (*Canis familiaris*) | (1.14)* |
| Cow (*Bos taurus*) | 1.06 |
| Common shrew (*Sorex araneus*) | 1.05 |
| Star-nosed mole (*Condylura cristata*) | 1.04 |
| Hispaniolan solenodon (*Solenodon paradoxus*) | 1.05 |

\* The weighted coverage of a genome to itself is parenthesized as it is not a comparative value

### Detection of single-copy orthologs

Single-copy orthologs (single gene copies or monoorthologs) are essential for the evolutionary analysis since they represent a useful conservative homologous set, unlike genes with multiple paralogs, which are difficult to compare between species. Longest proteins corresponding to each gene of *S. paradoxus* and three other *Eulipotyphla* – *Erinaceus europaeus*, *S. araneus*, *C. cristata* – were aligned to profile hidden Markov models of the TreeFam database [42,43] using HMMER [35]. Top hits from these alignments were extracted and used for assignment of corresponding proteins to families. The same procedure was performed in order to assign proteins to orthologous groups using profile HMMs of orthologous groups of the maNOG subset from the eggNOG database [44] as reference. Orthologous groups and families for which high levels of error rates were observed while testing assignment of proteins to them were discarded; the rest of the orthologous groups and families were retained for further analysis. Proteins and the corresponding assignments were obtained from the maNOG database for seven other species: *H. sapiens*,*M. musculus*, *B. taurus*, *C. familiaris*, *Equus caballus*, *Mustela putorius furo*, and *Monodelphis domestica*. Inspection of assignments across all the species yielded 4,416 orthologous groups containing single copy orthologous genes (**Database S5**).

### Species tree reconstruction and divergence time estimation

We used our genome assembly to infer phylogenetic relationships between *S. paradoxus* and other eutherian species with known genome sequences and estimated their divergence time using the new data. Based on the alignments of the single-copy orthologous proteins for the species included in the analysis, a maximum likelihood tree was built using RAxML [45] with the PROTGAMMAAUTO option and the JTT fitting model tested with 1,000 bootstrap replications. From the codon alignments of single-copy orthologs of the eleven species, 461,539 four-fold degenerate sites were extracted. The divergence time estimation was made by the MCMCtree tool from the software package PAML [46] with the HKY+G model of nucleotide substitutions and 2,200,000 generations of MCMC (of which the first 200,000 generations were discarded as burn-in). Divergence times were calibrated using fossil-based priors associated with mammalian evolution, listed in **Table 5** and based on [47–50]. FigTree [51] was used to plot the resulting tree, shown in **Figure 5**. According to this analysis, *S. paradoxus* diverged from other mammals of 73.6 Mya (95% confidence interval of 61.4-88.2 Mya). This is in accordance with earlier estimates based on nuclear and mitochondrial sequences (Roca et al., 2004; Brandt et al., 2016), and still within the timeframe of molecular estimates of divergence times between most island taxa and their mainland counterparts [52]. Our data supports solenodon divergence that occurred before divergence between shrews, moles and erinaceids [53–56], approximately at the same time as splits between other large mammalian groups, such as between rodents and primates, or carnivores and artiodactyls (**Figure 5**).

**Table 5.** Fossil-based priors associated with mammalian evolution used for calibration of divergence times [47–50]. The 4,416 single copy orthologs identified in our assembly were used for phylogeny inference via four-fold degenerate sites with programs RAxML [45] and PAML [46]. The resulting phylogenetic tree was plotted with FigTree[51] and is presented in **Figure 5**.

| Node | Calibration prior on clade | Node *min.* age (Mya) | Node *max.* age (Mya) | Evidence |
| --- | --- | --- | --- | --- |
| Opossum - placental mammals split | *Eutheria - Metatheria* | 157.3 | 169.6 | Fossil (Benton et al. 2015) |
| Human - mouse | *Archonta - Glires* | 61.5 | 100.5 | Biostratigraphy (Benton and Donoghue, 2007) |
| Primates, mouse - dog, horse, cow | *Euarchontaglires - Laurasiatheria* | 61.6 | 100.5 | Fossil (Benton et al. 2015) |
| Dog - ferret | *Canidae - Arctoidea* | 35 | 45 | Fossil (Wang et al., 2005; Munthe, 1998) |
| Solenodon - hedgehog, shrew, mole | *Lipotyphla* | 61.6 | 100.5 | Fossil (Benton et al. 2015) |
| Cow - horse | *Artiodactyla* as soft minimum | 52.4 | 100.5 | Fossil (Benton et al. 2015) |

**Figure 5.**
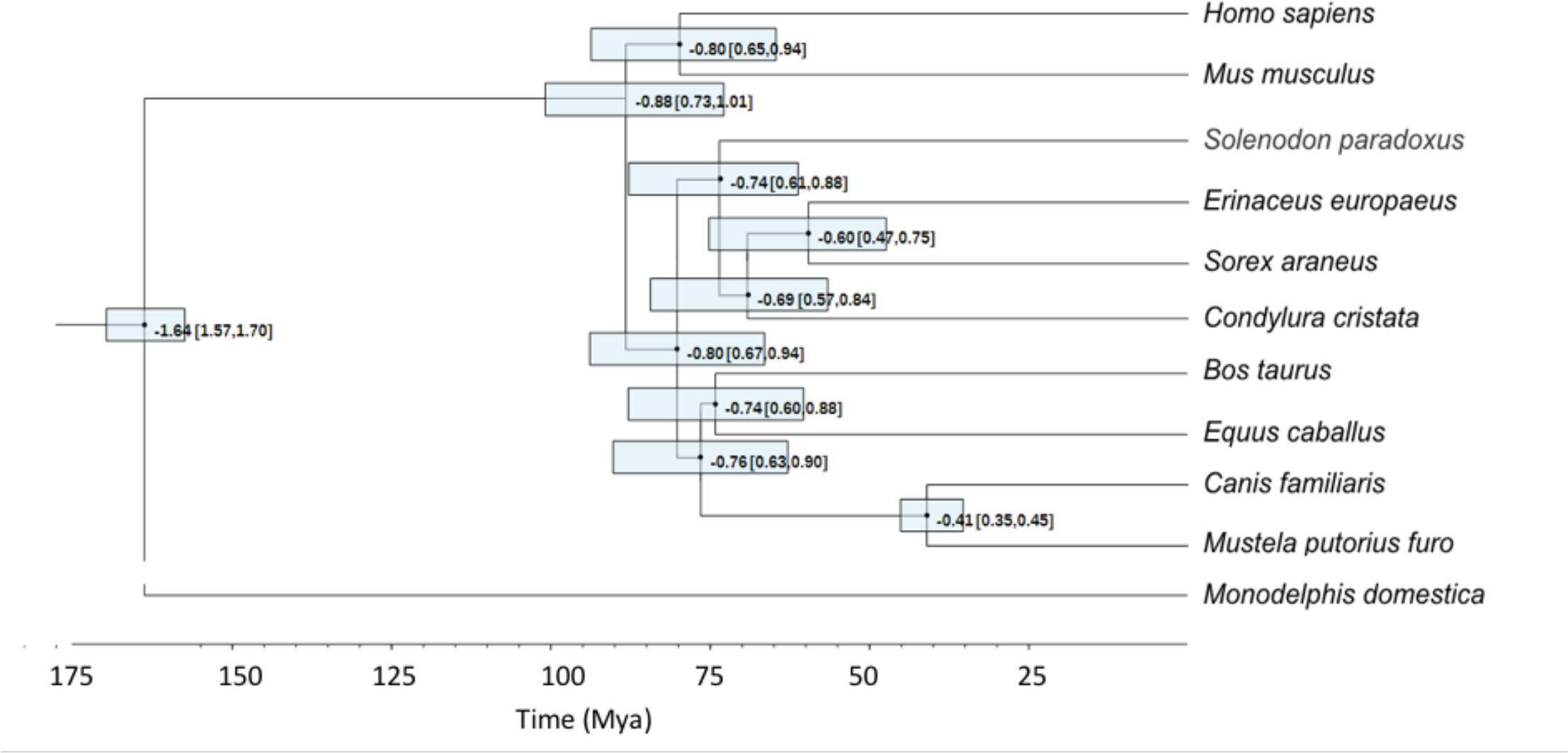
Divergence time estimates based on four-fold degenerate sites and on fossil-based priors (**Table 5**). The 95% confidence intervals are given in square brackets and depicted as semitransparent boxes around the nodes. The inferred divergence time of *S. paradoxus* from other mammals is 73.6 Mya (95% confidence interval of 61.4-88.2 Mya).

### Positively selected genes

To evaluate signatures of selection in the assembled genomes we used the dataset of the 4,416 orthologous groups containing single copy orthologous genes of the mammalian species described earlier. Single copy orthologs were used as a conservative set necessary to compare coding sequences that only arose one time in order to avoid using uncertainties associated with paralogs and lineage specific gene duplications. First, we translated DNA sequences into amino acids, aligned them in MUSCLE [57], and then translated back into DNA code using the original nucleotide sequences by PAL2NAL [58]. Genic dN/dS ratios were estimated among the 11 (including *Solenodon*) mammalian species used in constructing the phylogeny represented in **Figure 5**.

To estimate the dN/dS ratios, we used the *codeml* module from the PAML package [46]. The dN/dS ratios were calculated over the entire length of a protein coding gene. The branch-site model was not included in the current analysis because of the chance of reporting false positives due to sequencing and alignment errors [59], especially on smaller datasets, and additional uncertainties could be introduced from the lack of power under synonymous substitution saturation and high variation in the GC content [60].

All the single copy orthologs were plotted in the dN to dS coordinates and color-coded according to the 96 Gene Ontology generic categories (**Figure 6**). We retrieved values of dN, dS and w (w=dN/dS) for all single copy orthologs and used human annotation categories to assign all the genes with their gene ontologies (GO) using the Python package *goatools* [61] and the GO Slim generic database (GO Consortium, 2004) to assign the genes to the major GO categories.

**Figure 6.**
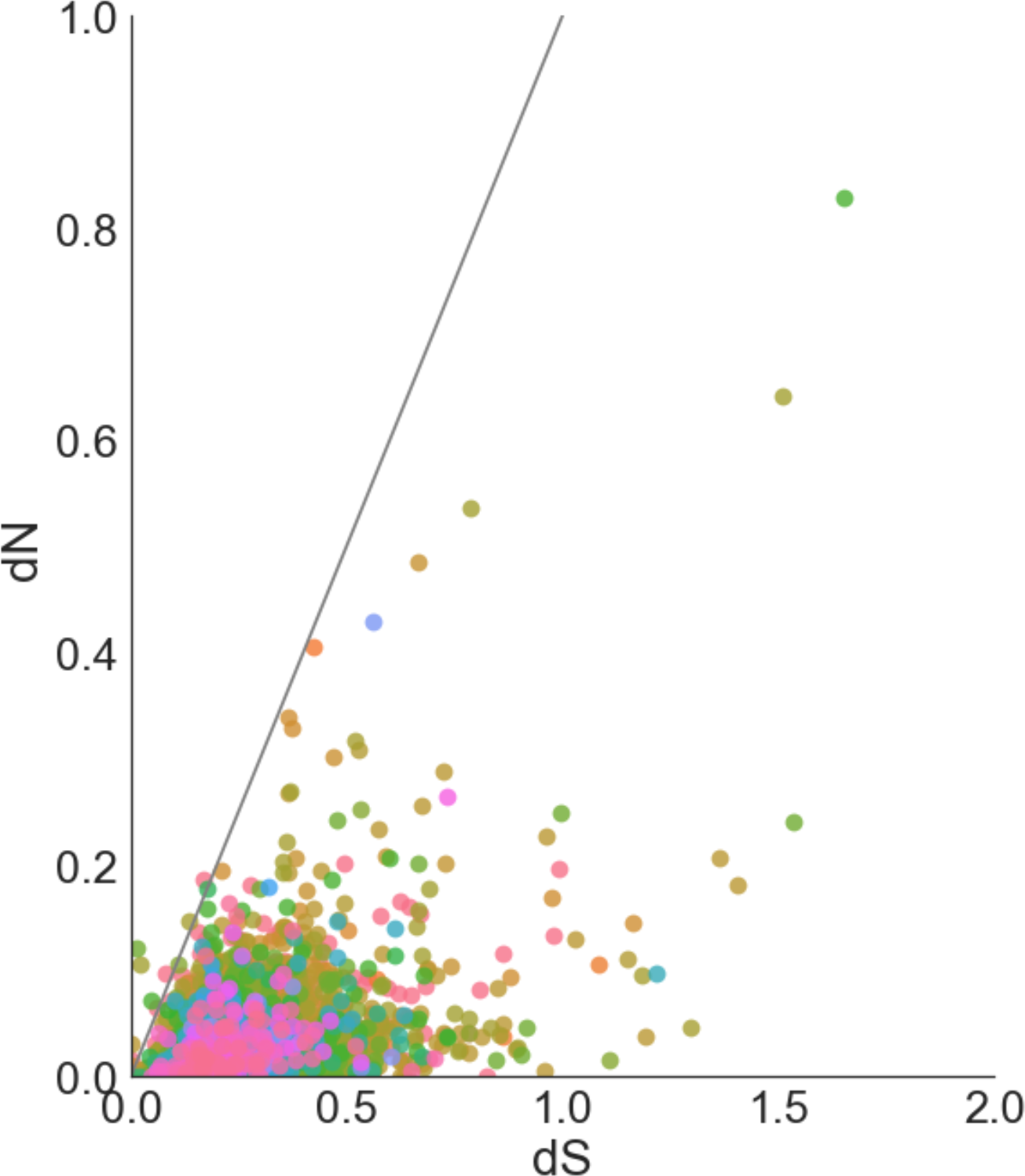
dN/dS ratios fort 4,416 orthologous groups containing single copy orthologous genes (monoorthologs). dN and dS ratios were calculated with the *codeml* module from the PAML package[46]. The dN/dS ratios were calculated over the entire length of a protein coding gene. Values are color-coded by GO term aggregated by the GO Slim generic database [61,62], and the color code legend is presented in **Figure S2**. The solid black line represents dN=dS; dots above it represent genes under positive selection. The figure is truncated at dN=1 and dS=2. All w, dN, and dS values are availablein **Database S6.**

The dN/dS values for the 12 genes exhibiting positive selection (**Table 6**) are visible above the dN=dS line. Three of these genes belong to the plasma membrane GO category (GO:0005886), while cytosol (GO:0005829), mitochondrial electron transport chain (GO:0005739), cytoplasm (GO:0005737) and generation of precursor metabolites (GO:0006091) were represented by one gene each. Five of the genes exhibiting positive selection signatures could not be assigned to the GO categories. Some of these are also associated with the plasma membranes (*TMEM56*, *SMIM3*), and one gene (*CCRNL4*) encodes a protein highly similar to the nocturnin, a gene identified as a circadian clock regulator in *Xenopus laevis* [63]. We are giving the full list of the genes, GO annotations, and the associated dN/dS values for each in the **Database S6**.

**Table 6.** The putative targets of positive selection in the Solenodon genome. The dN/dS values and the GO categories for the 12 genes that may be considered as candidates targets for positive selection in the *Solenodon paradoxus wodii* genome (dN>dS).All other genes are reported in **Database S6**.

| **Solenodon gene** | **dS** | **dN** | **dN/dS** | **GO category description** | **Human ortholog** |
| --- | --- | --- | --- | --- | --- |
| *ENOG410UG5H* | 0.000003 | 0.002563 | ≥999 | Plasma membrane | *KLF9* |
| *ENOG410USMX* | 0.000011 | 0.010830 | ≥999 | Plasma membrane | *TNFSF13B* |
| *ENOG410UWRE* | 0.000015 | 0.014790 | ≥999 | - | *SMIM3* |
| *ENOG410UNED* | 0.000174 | 0.030411 | 174.84 | - | *CCRN4L* |
| *ENOG410UJP8* | 0.013214 | 0.120449 | 9.12 | Cytosol | *PLK4* |
| *ENOG410UWA9* | 0.020955 | 0.104972 | 5.01 | Mitochondrion | *NDUFC1* |
| *ENOG410V3Q6* | 0.047538 | 0.071112 | 1.50 | Plasma membrane | *SYT16* |
| *ENOG410UQAM* | 0.078543 | 0.096445 | 1.23 | - | *WBP2NL* |
| *ENOG410UKXY* | 0.168982 | 0.185535 | 1.10 | - | *TIGIT* |
| *ENOG410UKXJ* | 0.134581 | 0.146926 | 1.09 | Cytoplasm | *LRRC66* |
| *ENOG410UIAB* | 0.060622 | 0.065402 | 1.08 | - | *TMEM56* |
| *ENOG410UG23* | 0.176172 | 0.177344 | 1.01 | Generation of precursor metabolites and energy | *THTPA* |

Traditionally, one of the most commonly used signatures of selection is expressed in terms of the ratio of non-synonymous (dN) to synonymous (dS) substitutions, dN/dS [64]. Synonymous rate (dS) expresses the rate of unconstrained, neutral evolution, so that when dN/dS<1, the usual interpretation is that negative selection has taken place on non-synonymous substitutions. Otherwise, when dN/dS>1, the interpretation is that the positive selection is likely to have accelerated the rate of fixation of non-synonymous substitutions. It is possible to quantify the proportion of non-synonymous substitutions that are slightly deleterious from the differences in dN/dS between rare and common alleles [65][66]. In our comparison, a subset of single copy orthologs dN/dS compared to the 10 mammalian species (**Figure 5**) is estimated to be ~0.18 or 18%, on average, compared for ~0.25 is reported for the human–chimp and ~0.13 reported for the mouse-rat comparisons [67]. In other words, it suggests that up to 82% of all amino acid replacements in *S. paradoxus* are removed by purifying selection [67].

Note that purifying selection is the conservative force in molecular evolution, whereas positive selection is the diversifying force that drives molecular adaptation. Overall the list of positively selected gene is relatively short compared to numbers of positively selected genes reported in other studies (e.g. human to chimpanzee comparison yields several hundreds of human-specific genes under selection [68–70]. This observation could be a consequence of the averaging effect of large comparison group that included mammals very distantly related to solenodons, but since it is expected that genetic drift was the principal driving force of the evolution in the solenodon genome over tens of millions of years of island isolation, a lower number of positively selected genes and predominance of the purifying selection is expected.

The dN/dS ratios can also be used as a proxy toto illustrate the rate of evolution for proteins. By looking at the trends in fast evolved genes (dN/dS > 0.25) we can make inferences about the factors that shaped the genome of this species during the millions of years of island isolation. To summarize the functional contributions, we used the PANTHER Overrepresentation Test and GO Ontology database based on the *H. sapiens* (**Table S1**) and *M. musculus* (**Table S2**) genes [71]. Interestingly, genes involved in the inflammatory response and located on cell surfaces were among those overrepresented among the rapidly evolving genes in Solenodon genome compared either to the human or mouse databases (**Table S1 and S2**).

### Venom gene identification

Since solenodon is one of very few venomous eutherian mammals, of special interest in the solenodon genome were the putative venom genes. While there was no saliva sample in our possession that could be analyzed for the expressed toxin genes, a comparative genome approach could be applied as an indirect way to find venom genes orthologous to genes expressed in venom for other species. First, we identified 6,534 toxin and venom protein representatives (Tox-Prot) from Uniprot [72], and queried them with BLAST against the current *S. paradoxus* genome assembly. The hit scaffolds were then extracted from the AUGUSTUS CDS prediction file. The same Tox-Prot sequences were used for Exonerate with the protein-to-genome model. The hits were used as queries against the NCBI database to ensure the gene identity, further validated through phylogenetic analyses with select model mammalian and venom reptilian genes (also adding randomly selected sequences for each gene, to reduce clade bias). The retrieved sequences were aligned with MUSCLE [57], followed by a Maximum likelihood (WAG+I+G) phylogenetic reconstruction. Hits were matched against their respective references in an alignment and visually inspected to assess potential venomous activity.

As a result, we identified 44 gene hits of the 16 most relevant protein venom classes (all present in snakes) in the *S. paradoxus* genome (**Table 7**). Inspection of pairwise MUSCLE alignments of the putative Solenodon venom genes with their animal homologs revealed several interesting cues. The putative venom genes could not be confirmed through genomic information alone, yet they cannot be discarded given that they were matched to high homology regions of closely related genes, such as those originally recruited into venom. There were also unusual insertions not found in other species’ venomous genes. Specifically, an insertion in a serine protease, a gene with a role in coagulation (namely coagulation factor X), is not present in known homologs. The insertion seems to be located at the start of the second exon. This particular gene was further analyzed to understand the insertion and its potential functional consequences (**Figure 7**). Finally, none of the venomous genes from the closest related venomous insectivore (*Blarina brevicauda*) have been found by this study. Our results indicate that a more detailed study of *Solenodon* venom genes using a transcriptome obtained from a fresh saliva sample is needed to address their molecular evolution and function.

**Table 7.** Homologous matches for the most relevant protein venom classes in the *Solenodon paradoxus* genome. Genes were identified by querying 6,534 toxin and venom protein representatives found in animal venoms in Tox-Prot from Uniprot [72]. All of the protein groups are present in snake venoms. The sequences of the putativevenom genes from *S. paradoxus* are available in the **Database S7**.

| Protein groups<br>found in animal venoms | Number of hits in the<br>*S. paradoxus* genome |
| --- | --- |
| Metalloproteinase; Serine protease | 8 each |
| Hyaluronidase | 6 |
| (Acetyl)Cholinesterase | 2 |
| Calglandulin; Nerve growth factors | 4 each |
| Lipase | 3 |
| Hydrolase; Kunitz serine protease inhibitor; Nucleotidase; O-methyltransferase; Oxidase; Peptidase; Phosphodiesterase; Phospholipase; Vascular endothelial growth factor | 1 each |

**Figure 7.**
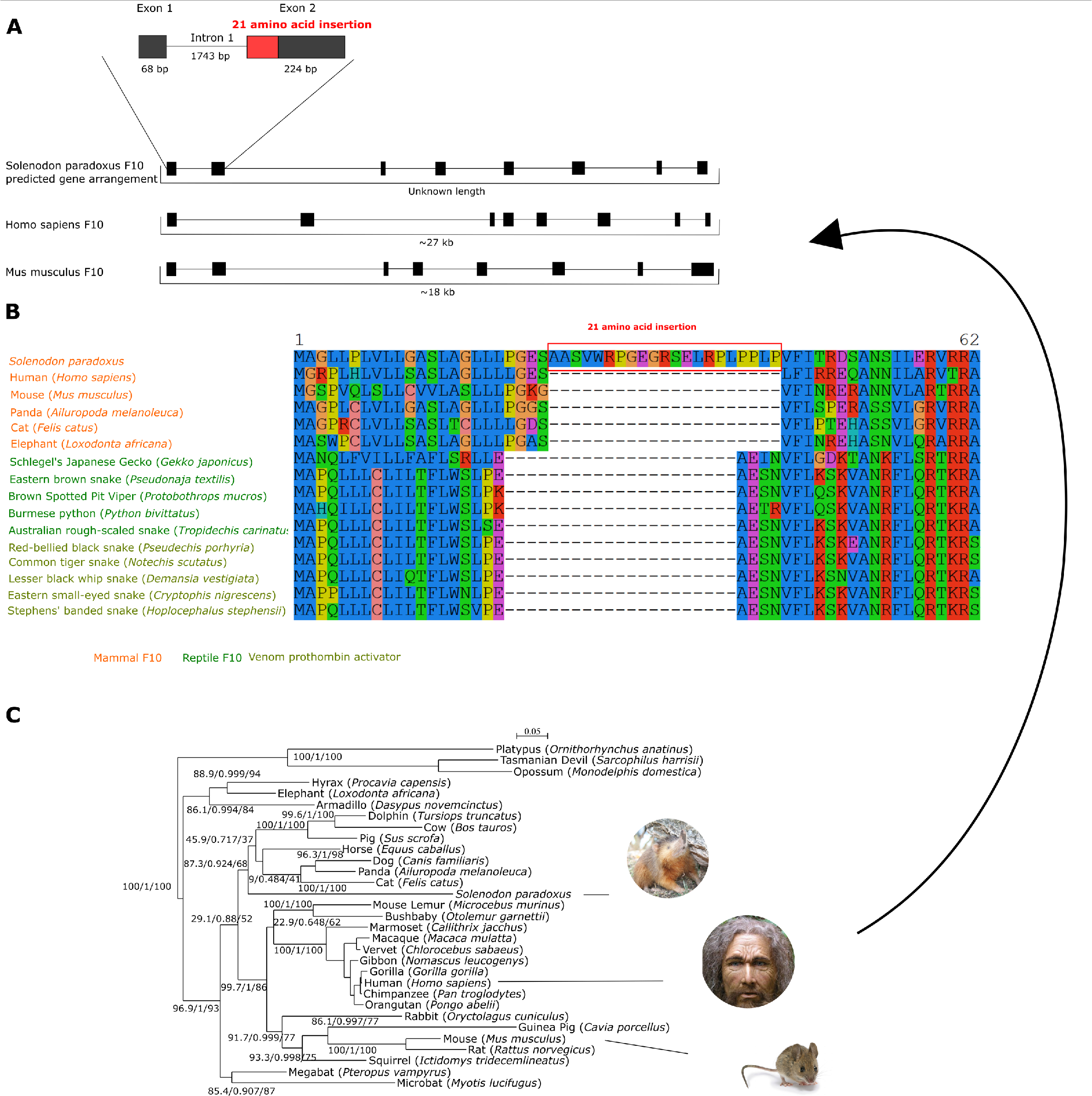
(A) Predicted coagulation factor X (F10) gene structure arrangement from known homologs organization (due to the scaffolding, the total gene length is unknown in solenodon). The 21 amino acids insertion is highlighted in red on the exon two of the solenodon F10 gene. Exons are represented as black boxes and introns as lines connecting exons. (B) F10 protein sequence alignment showing an unusual insertion in the *Solenodon paradoxus* genome absent in all other mammalian and reptilian genes retrieved from the Tox-Prot from Uniprot [72]. The insertion of 21 amino acids is indicated with a red-boxed line in the alignment. (C) Reconstructed mammal F10 phylogenetic tree using Maximum-likelihood model GTR+I+Γ, 1000 bootstraps (1590 bp-long alignment). The numbers set indicate approximate likelihood-ratio branch test (aLRT), Bayesian like modification of the aLRT and bootstrap percentage, respectively.

### Genomic variation and demographic history inference

Once the reference alignment was assembled as a consensus between the sequences obtained from the five *S. p. woodi* individuals, polymorphisms were identified in the six individual genomes by aligning them to the combined reference. Single-nucleotide and short variants and indels were identified in five southern and one northern individual using Bowtie2 [73], SAMtools and Bcftools [74], and VCFtools [75]. Each of the *S. p. woodi* individuals differed from the reference by an average of 1.25 million polymorphisms, and the *S. p. paradoxus* individual differed by 2.65 million from the reference assembly.

Whole genome SNV rates for solenodon were calculated, defined as a ratio of all observed SNVs to all possible SNV sites in the genome were found to be comparatively low among other mammals (**Figure 8**) [76–79]. To enable this comparison, the same calculations were employed, where SNVs were not filtered by repetitive regions or mappability mask and the number of possible SNV sites was defined as the genome assembly size minus number of unknown base pairs (’N’).

**Figure 8.**
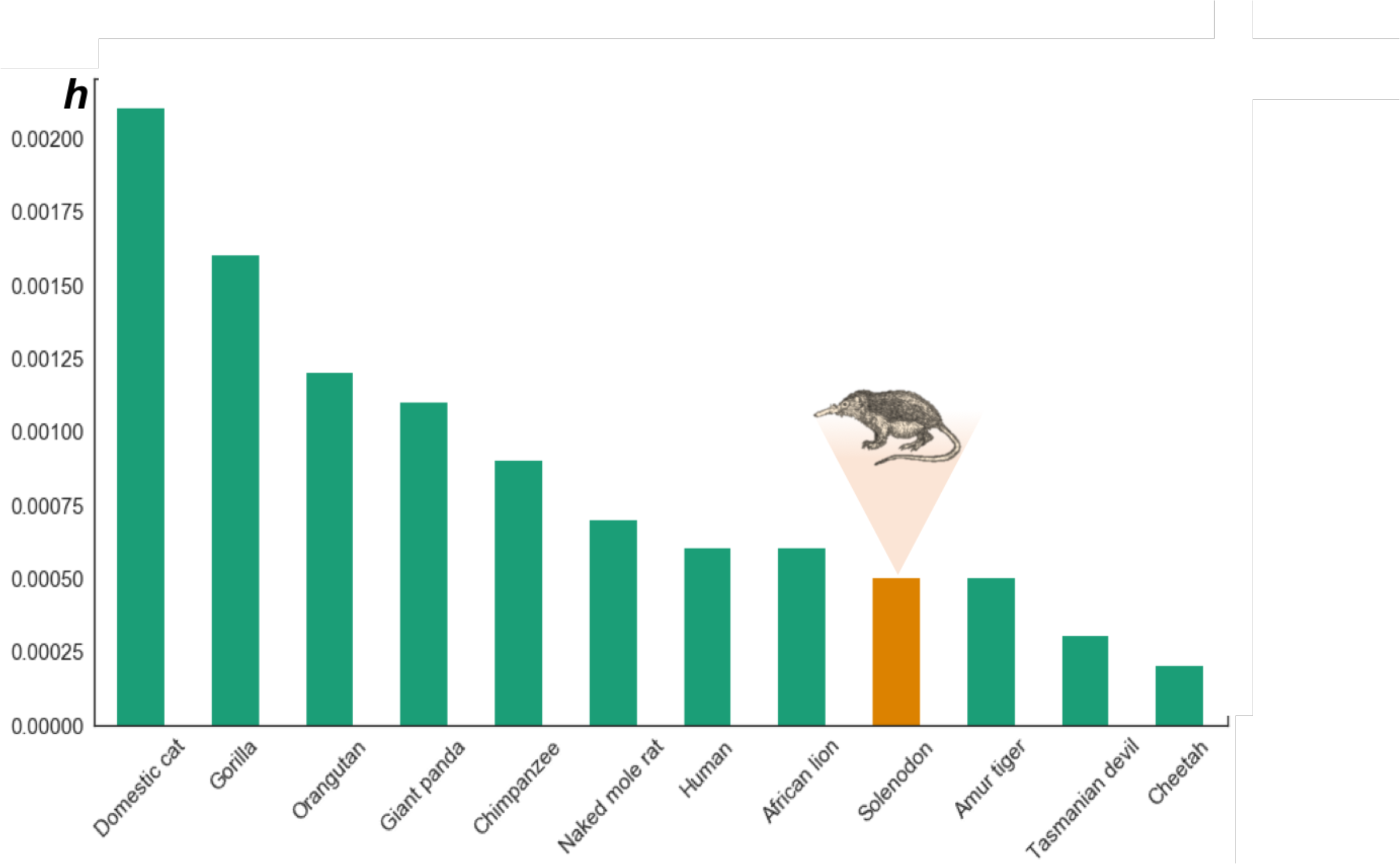
Low genome heterozygosity in *Solenodon paradoxus* compared to other mammalian species. The SNV rate in the *S. paradoxus woodii* genome is shown relative to other mammal genomes as an estimate of genome diversity (*h*). The rate for each sequenced individual was estimated using all variant positions, with repetitive regions not filtered. The SNVs are deposited in the **Database S9**.

Based on the variation data from the genomes of two subspecies (*S. p. woodi and S. p. paradoxus*), we estimated population dynamics using Pairwise Sequentially Markovian Coalescent (PSMC) model [80]. PSMC uses the coalescent approach to estimate changes in population size: since each genome is a collection of hundreds of thousands independent loci, it allowed us to create a TMRCA distribution across the genome and estimate the effective population size (*Ne*) in recent evolutionary history (e.g. from 10,000 to 1 million years).

Demographic history was inferred separately for *S. p. woodi and S. p. paradoxus*, and the resulting plots revealed differencesa difference in demographic histories of the two subspecies (**Figure 9**). Each southern individual was considered separately and their demographic histories are overlaid. The difference in demographic history provides another argument in favor of a subspecies split, as evidenced by distinctly different effective population sizes at least since 300 Kya. According to this analysis, the northern solenodon subspecies currently has a much larger *Ne*, which has expanded relatively recently, between 10,000 – 11,000 years ago (**Figure 9**). Prior to that, it was the southern subspecies (*S. p. woodi*) who had a larger *Ne*. At the same time, the demographic history inference for both populations show similar cyclical patterns of expansion and contraction around the mean of 6,000 individuals for the southern subspecies (*S. p. woodi*) and 3,000 for the northern subspecies (*S. p. paradoxus*).

**Figure 9.**
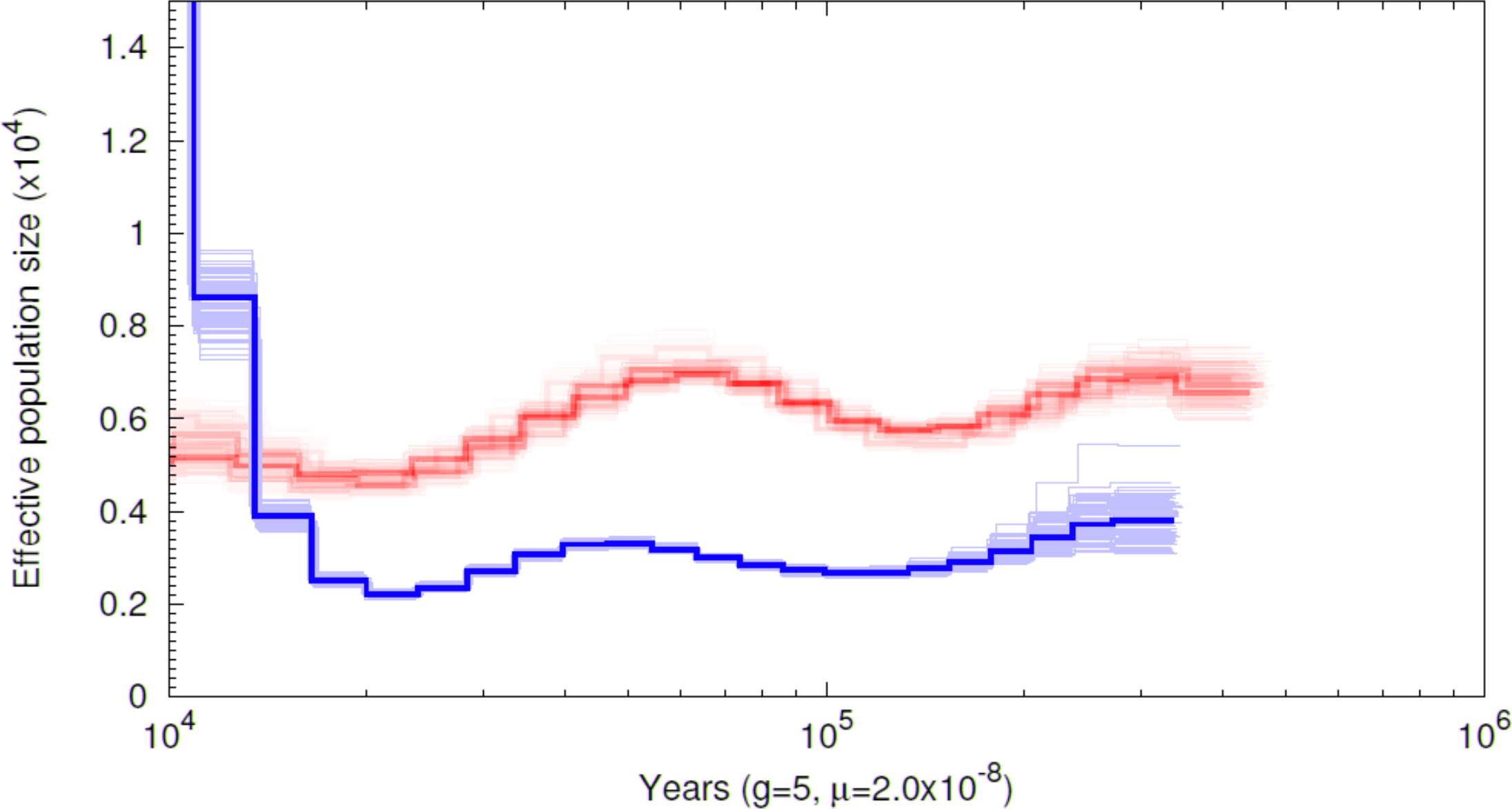
Demographic history inference for the southern (red) *S. p. woodi* and the northern (blue) *S. p. paradoxus* subspecies using the pairwise sequentially Markovian coalescent (PSMC) model [80].

### Development of tools to study population and conservation genetics of S. paradoxus

The presence of genome wide sequences of multiple individuals from two subspecies created a possibility for the development of practical tools for conservation genetics of this critically endangered species. Generally, microsatellite loci are both abundant and widely distributed throughout the genome sequence, and each locus is characterized by a unique flanking DNA sequence so it can be independently amplified in many individuals [81–83]. The major advantages of microsatellite markers are well known: codominant transmission, high levels of polymorphisms leading to the high information content and higher mutation rate that allows differentiation between individuals in the same population. Finally, microsatellite markers are easy to genotype even with the most basic laboratory configurations. While a genome obtained from one individual can be searched for the potentially variable microsatellite loci, it would (1) miss the majority of loci not represented in the individual’s two chromosome sets, and (2) result in many false positives that must be verified by laboratory tests (usually by electrophoresis of the amplified fragments from population samples). Therefore, the availability of several genomes would contribute to (1) more comprehensive set of variable markers, and (2) these markers would be more likely to be true positives with much higher probabilities to show variation between individuals.

All three assemblies from this study (A, B and C) were independently analyzed using a Short Tandem Repeat (STR) detection pipeline. A, B and C assemblies were analyzed separately with TRF (Tandem Repeats Finder) to locate and display tandem repeats [84]. Each of the six individual samples from the two Solenodon subspecies (five from *S. p. woodi* and one from *S. p. paradoxus*) were aligned to the reference assemblies A, B, and C by Burrow-Wheelers Aligner (Li and Durbin, 2009). Each set of individual alignments was analyzed with HipSTR [85]. Only the loci that shared more than 20 reads in the provided samples alignments were considered forfurther variable microsatellite loci search. The result of this search was saved in a Variant Call Format (VCF) file that included annotations of all loci that had variation between the samples and passed the minimum qualification of the reads parameter: to be successfully genotyped in at least one sample. The loci that did not pass these criteria were labelled as “*unsuccessfully verified*“ and excluded from the list.

The remaining loci were subjected to additional filtering: all genotypes that had less than 90% posterior probability according to HipSTR [85], genotypes with a flank indel in more than 15% of reads, and genotypes with more than 15% of reads with detected PCR stutter artifacts were discarded. The final set contains loci that have at least two allele calls between the individuals after filtering have been deposited in the polymorphic microsatellite database (**Database S8**). This database contains a list of variable microsatellites discovered, the 1200 bp flanking sequence for primer construction, and the information on whether and where it was found variable - between subspecies, or inside one of the subspecies. We also report the type (di-tri-, etc), number of repeats, number of variants, % variable, and provide up to 100bp flanking sequence on both sides that can be used to develop primer sequences (**Database S8**).

## Discussion

In this study, we sequenced and assembled the genome of an endangered Caribbean mammal that survived tens of millions of years of island isolation, but nevertheless is currently threatened by extinction due to anthropogenic activities. Our approach demonstrated sequencing, assembly and annotation of a genome of one of the earliest branches that split from the placental mammal tree, and provides insight into the de novo sequencing of other challenging genomes by delivering an important phylogenetic reference to the mammalian evolutionary history which can be added to the growing list of other phylogenetically diverse mammalian genomes for analysis in a comparative context [86]. Albeit the full description of genome diversity of this rare enigmatic mammal needs to be further improved with more samples and analyses, our initial assembly of the solenodon genome contributes information and tools for future studies of evolution and conservation. Future studies can combine the current genome annotations with the inclusion of additional genetic and ecological data from further sampling.

With the new genome-wide assembly, we produce a phylogeny that validates previous estimates of the time of the Solenodon divergence from other eutherian mammals [3,13]. Our comparative genome analyses have facilitated investigation of the timing of divergence of solenodon, and provide a window into genetic underpinnings of adaptive features, making it possible to begin to investigate the phenotypic characteristics of these unique animals including genes responsible for inflammation and venoms, and how these may reflect its adaptation. In addition, we developed tools that will help guide the future genome studies as well as conservation surveys of the remaining solenodon populations on the island of Hispaniola. In this study, we have made the first step into the whole-genome analysis of the *Solenodon*. A more complete genome sequence may provide a better picture of its evolutionary history, possible signatures of selection, and clues about the genetic basis of adaptive phenotypic features facilitating life in the Caribbean islands, and contribute to a better insight of island evolution and possible responses to current and future climatic changes.

### The string graph assembly approach for homozygous genomes

The advantages of the string graph assemblies in our particular case can be understood by looking at the nature of the underlying algorithms. The de Bruijn graph is a mathematical concept that simplifies genome assembly by reducing information from short next generation sequencing reads, of which there can be billions, to an optimized computational problem that can be solved efficiently [87]. However, some information may indeed be lost, as the set of reads is effectively replaced with a set of much shorter k-mers to produce an optimal assembly path. Usually, this is compensated by overwhelming amounts of data in high coverage assemblies, and the difference in effectiveness between this and other types of algorithms, barring speed, becomes less evident. While sequencing becomes cheaper, genome projects continue to rely on the increased high quality coverage, increasing the cost of the sequence data rather than trying to increase the efficacy of the assembly itself. In contrast, the string graph-based algorithms for genome assembly are intrinsically less erroneous than de Bruijn graph based ones, since building and resolving a string graph does not require breaking reads into k-mers and therefore does not sacrifice long-range information [17]. This also helps reduce the probability of mis-assemblies: in theory, any path in a string graph represents a valid assembly [88,89]. String graph based approaches have already been applied successfully to assemblies from high coverage read sets; and a one example is the Assemblathon 2 [90]. In projects with lower genome coverage like ours, adoption of string graph based approach might be of benefit to the genome assembly because it uses more information from the sequences. However, there are two major downsides for the widespread use: (1) it is more computationally intensive than methods utilizing de Bruijn graph algorithms, and (2) the implementation of the string graph model is sensitive to sequence variation, and the effectiveness of this approach may depend on the level of heterozygosity in a DNA sample. It is worth noting that Fermi [17] was primarily intended for variant annotation via *de novo* local assembly, and not for whole genome assembly. Nevertheless, the new genome-wide data produced by our pipeline was sufficient for the comparative analysis, and has been annotated for the genes and repetitive elements, and interrogated for phylogeny, demographic history and signatures of selection. In addition, using the current genome assembly we were able to annotate large transpositions and translocations in the *Solenodon* in relation to the closest available high-quality genome assembly (*S. araneus*).

## Potential implications

### Comparative genomics

We have taken advantage of the fact that the genome of this mammal is extremely homozygous, which allowed us to combine samples of multiple individuals in order to provide higher coverage and achieve a better assembly using Illumina reads. The current assembly was performed without the use of mate pair libraries and high quality DNA, nevertheless it is comparable in quality to other available mammalian assemblies. In terms of contig N50 as a measure of contiguity, our assembly resulted in contig N50 of 54,944 while the most closely related available genome sequences of *Sorex araneus* (SorAra2.0)assembly features a contig N50 of 22,623, and the *Condylura cristata* (ConCri1.0) assembly has contig N50 of 46,163. It should be noted that scaffold N50 values are not to be compared as this study used only paired end reads, as opposed to *S. araneus* and *C. cristata*. More importantly, the assembly provided annotation for more than 95% of the genes and allowed the subsequent comparative analysis.

Specifically, the repetitive composition of the solenodon genome was evaluated. Compared to the estimates based on the reference human genome [91], very conspicuous is the lower numbers of SINEs (no *Alu* elements), and a significantly lower number of LINEs as well. Transpositions and translocations between the genomes of *S. paradoxus* and *S. araneus* were identified; very few rearrangements and translocations between the assembly and the *S. araneus* genome were found. At the same time a higher coverage would be needed to do more detailed analyses, for instance to address the relative length and similarity of indels and copy number polymorphisms between solenodon populations [92].

### Evolutionary genomics

As a result of the additional information for the nuclear genomes, we were able to confirm earlier divergence time estimates based on a set of genes [3], as well as full mitochondrial sequences [13]. The whole genome analysis points to a split between solenodon and the insectivores that occurred around 74 Mya (**Figure 5**), which is very close to our earlier estimates of 78 Mya, based on the full mitochondrial genome [13]. Our result does not support the 60 Mya estimate made by the phylogenetic analysis based on sequences of five slowly evolved nuclear genes [15].

Our assembly provided enough gene sequences to gain an insight into the evolution of functional elements in the solenodon genome. It is reasonable to suggest that this species historically had low effective population sizes, if they remained close to those estimated by this study: or about 4,000 on average (**Figure 9**). Genetic drift is the prevailing force in small populations, so we did not expect to see many signatures of positive selection. Nevertheless, among the 4,416 single copy orthologs analyzed for dN/dS ratios over the entire length of a protein-coding gene between *S. paradoxus* and 10 other mammals, 12 genes were identified as positively selected. Among these, the majority were membrane proteins, and one gene (*CCRNL4*) a possible circadian clock regulator (**Table 6**). It is possible that the short list of the positively selected gene could be a consequence of the large comparison group that included mammals very distantly related to solenodon, and its genes need to be compared with more closely related species, for example once the genome of *S. cubanus* is reported, and better gene annotations for *Sorex araneus* become available.

*Solenodon* is one of few mammals that use venomous saliva to disable prey, but is unique because it delivers its poison similarly to snakes — using its teeth to inject venomous saliva into its target. Different approaches could be used to characterize venom genes, such as the use of non-curated databases to widen the search spectrum thatwhich may include some potentially different molecules that could be found in Solenodon. For example, 6,534 toxin and venom protein representatives can be found in the UniProt database. It is also important to note that many venomous sequences currently found in databases may not match *Solenodon*’s particular genes given the species’ deep divergence from any other related venomous mammalian species. The fact that hits to known curated venoms were not fully determined suggests that the *Solenodon*’s venom may contain novel protein modifications with unknown potential or application, making it valuable for future detailed characterization.

Genes with hits to venom sequences, such as serine proteases involved in coagulation (namely the coagulation factor X) are of major interest, since these genes in solenodon exhibited unusual insertions when compared to their homologs (**Figure 7**). The detection of an unusual insertion in serine proteases has been previously found in another venomous mammalian species, the shrew *Blarina brevicauda*, but in solenodon occurs in both a different gene and site. This particular gene from solenodon, the coagulation factor X, is involved in the circulatory system and responsible for activating thrombin and inducing clotting. The insertion in the coagulation factor X gene seems to be a hydrophilic alpha helix with three potential protein-protein interaction sites. It occurs at the end of the region annotated as the signal peptide, while having a signal peptide cleavage site itself at the beginning of its sequence. The factor X protein structure was successfully modeled by Swiss-Model based on the venomous elapid snake *Pseudonaja textilis* (pdb: 4bxs), to have a heavy chain that contains the serine protease activity, which was modeled with a high degree of confidence (**Figure 10**). The venom prothrombin activator has an advantage as a toxin in part due to modifications in inhibition sites, making it difficult to stop its activity. Another advantage is that the molecules are always found in an active form (Kinin). We hypothesize that the insertion could allow a more successful interaction with molecules capable of activating the F10 protein. Both *Solenodon’s* extract and venom prothombin activator injections in mice can be lethal in minutes [7,93]. The insertion was also searched against possible mobile DNA elements, but no matches were found. Our advanced results should be followed in the future by detailed pharmacological studies.

**Figure 10.**
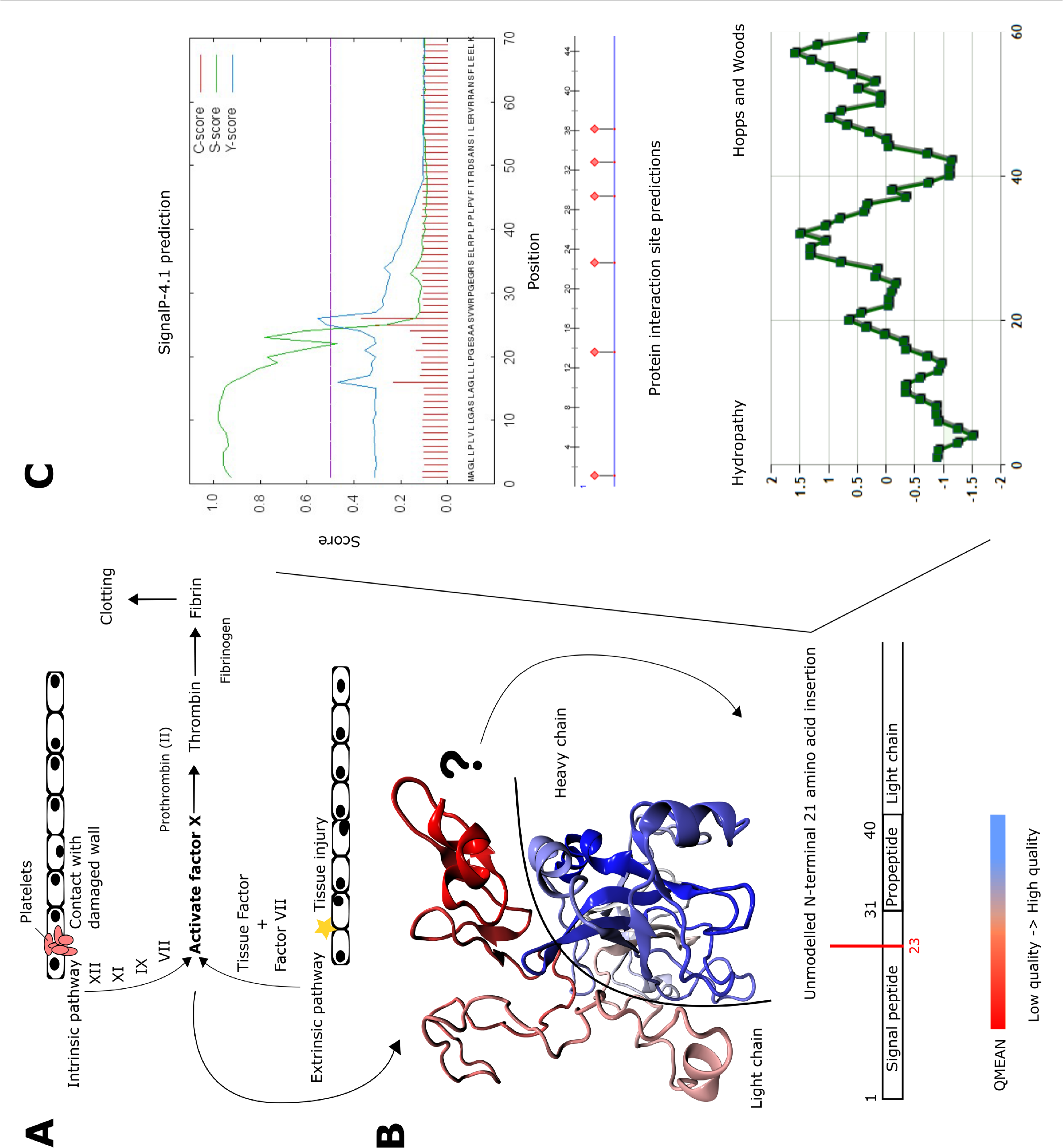
(A) Simplified version of the coagulation cascade, indicating key steps involving the coagulation factor X (F10). (B) Protein modeling of solenodon’s sequence data using SWISS MODEL. The target model (4bxs) used was the venomous elapid snake *Pseudonaja textilis*’ F10 like protease. Due to its location the insertion cannot be represented in the model (its location is indicated according to the PDB annotation). Colors indicate model quality, with red being low quality and blue high quality modeling. Colors also separate F10’s light chain (EGF-like domain) in red from the heavy chain (serine protease domain) in blue (the half circle line in black separates both domains). (C) Amino acid sequence properties calculated for the solenodon’s F10 translated gene, with focus on the insertion region 23-43. One signal peptide cleavage site was detected between position 25 and 26. Predicted protein interaction sites at position 26, 29-30 and 32-40. Hydropathy analysis showed the relatively hydrophilic structure for the insertion.

### Conservation genetics

The low variation that exists between the solenodon sequences is hardly surprising, because the theoretical consensus in conservation genetics predicts that populations with a smaller *Ne* lose genetic diversity more rapidly than populations with a larger *Ne* [94], and measures of genetic diversity have been explicitly suggested to IUCN as a factor to consider in identifying species of conservation concern [95]. The low *N_e_* in each subspecies is confirmed by our analysis (**Figure 9**), and shows particularly low levels in *S. p. woodi*. Due to the limitations of PSMC, the most recent *Ne* cannot be calculated from the genome sequences [80]. Therefore, this level of diversity indicates historic levels persisting for at least 120,000 years, and does not reflect the recent impact on the solenodon population caused by anthropogenic factors in the last 10,000 years (**Figure 9**).

Many endangered species with small populations also have reduced heterozygosity levels across their genomes, and would benefit from a computational approach that reduces the cost and optimizes the amount of data for the genome assembly. The real-life scenarios where no high-quality DNA can be produced because of the remoteness of sampling location, difficulty in transportation and storage, or when the high coverage cannot be produced due to the limited funds are well known to many, especially in the field of conservation genetics. The difficult field conditions and international regulations make it difficult to obtain samples with high molecular weight DNA. To aid the future conservation studies, we intend to mine the current dataset for microsatellite markers that can be used for the identification of subspecies, and potentially the populations of solenodons, as well as to be used as tools for studies on population diversity and monitoring.

The comparative analysis of the number and the length of microsatellite alleles pointed once more to the advantage of assembly B over A and C. The average length of microsatellites in assembly B is the highest (20.95 (assembly A), vs. 21.14 (assembly B) vs. 18.86 (assembly C)), which also indicated the advantage of assembly B over the alternatives (A and C). This may be the direct consequence of the higher number of microsatellite allelesthat were successfully genotyped in all of the southern samples for assembly B (2,660), as well as the number of variable microsatellites detected and variable but fixedvariability between the two subspecies (639). The large drop in the number of variable microsatellites between the two categories may be explained by the reduced amount of information that can be obtained from the single genome of the northern subspecies (*S. p. paradoxus*) in this study. However, there are 170 variable microsatellites found exclusively in the northern subspecies, which can be used as ancestry markers to evaluate population structure and migration rates between the subspecies. The Venn diagrams representing microsatellite variation in three assemblies are presented in **Figure 11.**

**Figure 11.**
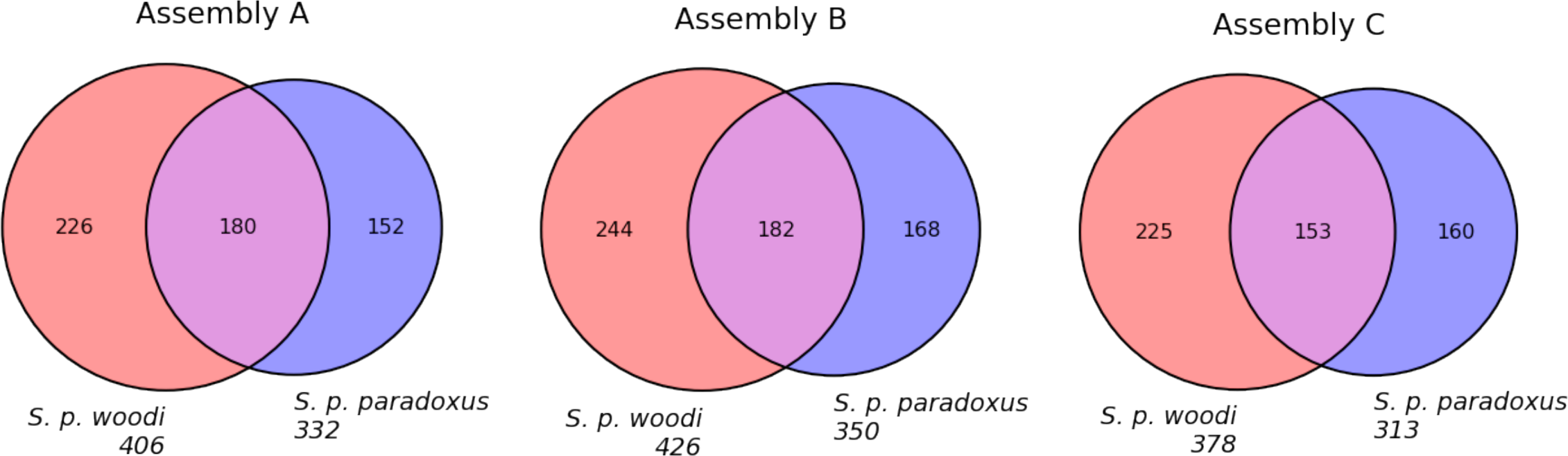
Numbers of variable microsatellite alleles discovered in *S. paradoxus* assemblies. The diagrams were built independently for Fermi-based assemblies (A and B) and one SOAPdenovo2 based assembly (C). The red circle indicates microsatellites that were successfully genotyped in all samples with at least one alternative allele in the southern subspecies (*S. p. woodi*). The blue circle indicates microsatellites that were successfully genotyped in all samples with at least one alternative allele in the northern subspecies (*S. p. paradoxus*). The overlap indicates microsatellite loci with at least one alternative variant found in both subspecies. All alleles discovered, number of fixed alleles in each population and number of unique alleles in each population are presented in **Table S3.** All the candidate microsatellite loci discovered in this study, along with their 5’ and 3’ flanking regions are listed in the **Database S8.**

Finally, new data confirms the north–south subspecies subdivision within *S. paradoxus* reported earlier (Brandt et al. 2016). Moreover, the southern Hispaniolan solenodons currently have a much smaller *Ne*, which are recently expanded based on the pairwise sequentially Markovian coalescent (PSMC) model [80]. Moreover, according to our analysis, this difference in *Ne* between the two subspecies has existed for at least 120 thousand years (**Figure 9**). This separation has been suggested by the earlier study using full mitochondrial DNA (Brandt et al., 2016). Recently, another genetic survey using mitochondrial cytochrome b and control region sequences from 34 solenodon samples identified unique haplotypes in each biogeographic region [14]. The island of Hispaniola has been historically divided into three main biogeographic regions that differ in climate and habitat. The north and center of the island provide the largest area with known solenodon populations, and shows no discontinuity with the southeast. However, the solenodon populations in the southwestern part of the island are currently geographically isolated by Cordillera Central, and may have been isolated in the past by the ancient island divide across the Neiba Valley (**Figure 2**). This geographic isolation is likely the reason why the *S. p. paradoxus* in the larger northern area, and *S. p. woodi* in the southwest, show morphological differences suggestive of separate subspecies [12]. Future conservation strategies directed at protecting and restoring solenodon populations on Hispaniola should take into consideration this subdivision, and treat the two subspecies as two separate conservation units. Unfortunately, we did not have a chance to confirm the identity of a small remnant population that also survives at the Massif de la Hotte in the extreme western tip of Haiti [14,96], and, if procured, may show genetic divergence from the two populations described in our study.

## Methods

Locations of where the samples were obtained are described on the map (**Figure 2**), and coordinates are listed in **Table S4**. Solenodons were caught with help of local guides (Nicolás Corona and Yimell Corona). Durisopropyl alcohol. Once collecteding the day, potential locations were inspected in daylight for animal tracks, burrows, droppings and other signs of solenodon activity. At dawn, ambushes were set up in the forested areas along the potential animal trails. The approaching solenodons were identified by sound, and chased with flashlights when approached. Since solenodons move slowly, animals were picked up by their tails, which is the only way to avoid potentially venomous bites. All wild caught animals were released back into their habitats within 10 minutes after their capture. Before the release, the animals’ tails were marked with a Sharpie pen to avoid recapturing.

Blood was drawn by a licensed ZooDom veterinarian (Adrell Núñez) from the *vena jugularis* using a 3mL syringe with a 23G x 1” needle. The blood volume collected never exceeded 1% of body weight of animals. Before the draw, an aseptic technique was applied using a povidone–iodine solution, followed by isopropyl alcohol. Once collected, the samples were transferred toa collection tube with anticoagulant (BD Microtainer, 1.0mg K2EDTA for 250–500lL volume). Collection tubes were refrigerated and transported to the lab at the *Instituto Tecnológico de Santo Domingo* (INTEC) where DNA was extracted from samples using the DNeasy Blood & Tissue kit (Qiagen, Hilden, Germany). This study has been reviewed and approved by the Institutional Animal Care and Use Committee of the University of Puerto Rico at Mayagüez (UPR-M). All the required collection and export permits issued by the US government under the Endangered Species Act (ESA), Convention on International Trade in Endangered Species of Wild Fauna and Flora (CITES), by the Animal and Plant Health Inspection Service (APHIS) and the Ministry of the Environment and Natural Resources of the Government of the Dominican Republic had been obtained before any field work was started.

### Sequencing

Sequences were generated by Illumina HiSeq (Illumina Inc). The Illumina HiSeq generated raw images utilizing HCS (HiSeq Control Software v2.2.38) for system control and base calling through an integrated primary analysis software called RTA (Real Time Analysis. v1.18.61.0). The BCL (base calls) binaries were converted into FASTQ utilizing the Illumina package bcl2fastq (v1.8.4). The sequencing data for each sample used in this study is presented in **Table S5**.

## Availability of supporting data and materials

**Database S1:** Lists of repeats for of in the Solenodon genome (assemblies A and B) http://public.dobzhanskycenter.ru/solenodon/repeats/solpar-a.txt http://public.dobzhanskycenter.ru/solenodon/repeats/solpar-b.txt

**Database S2:** List of protein coding genes in the Solenodon genome (assembly B) http://public.dobzhanskycenter.ru/solenodon/genes/solpar-b.gff also cds for each gene and translated sequences

**Database S3:**List of the annotated non-coding RNAs in the Solenodon genome http://public.dobzhanskycenter.ru/solenodon/rna

**Database S5:** List of monoorthologs in the Solenodon genome (columns include: ENOG id, gene name) http://public.dobzhanskycenter.ru/solenodon/monoorthologs.txt

**Database S6:** List of genes with dN/dS values and GO annotations http://public.dobzhanskycenter.ru/solenodon/selection.xls

**Database S7:** List of venom genes http://public.dobzhanskycenter.ru/solenodon/venom_genes_HitGeneDB.fasta

**Datablase S8:** Microsatellite loci discovered in genomes of two solenodon *subspecies Solenodon paradoxus paradoxus* (northern) and *S. p. woodi* (southern), alleles, flanking regions (1200 bp), and frequency information for the two subspecies http://public.dobzhanskycenter.ru/solenodon/STRs.xlsx

**Database S9:** Lists of single nucleotide differences (SND) from the assembled individual genome (Spa-1 (from *Solenodon paradoxus paradoxus*) and Spa K, - L, - M, -N, and –O (from the five *S. p. woddii*)) used to show estimate of heterozygosity in Figure *(see explanation in text) http://public.dobzhanskycenter.ru/solenodon/variants

## Declarations

Authors have declared that they do not have any competing interests

## Endnotes

The authors thank Nicolás Corona and Yimell Corona for assistance in collecting samples. This research was supported in part by NSF award #1432092.

